# Neurological mechanism of sensory deficits after developmental exposure to non-dioxin-like polychlorinated biphenyls (PCBs)

**DOI:** 10.1101/2021.02.14.430009

**Authors:** Nadja R. Brun, Jennifer M. Panlilio, Kun Zhang, Yanbin Zhao, Evgeny Ivashkin, John J. Stegeman, Jared V. Goldstone

## Abstract

The most abundant polychlorinated biphenyl (PCB) congeners found in the environment and in humans are neurotoxic. This is of particular concern for early life stages because the exposure of the more vulnerable developing nervous system to chemicals can result in neurobehavioral disorders. To uncover currently unknown links between PCB target mechanisms and neurobehavioral deficits, we investigated the effects of the non-dioxin-like (NDL) congener PCB153 on neuronal morphology and synaptic transmission linked to the proper execution of a sensorimotor response using zebrafish as a vertebrate model. Zebrafish that were exposed during development to concentrations similar to those found in human cord blood and PCB contaminated sites showed a delay in startle response. A similar delay was observed for other NDL congeners but not for the potent dioxin-like congener PCB126. Morphological and biochemical data demonstrate that while exposure to PCB153 induced swelling of afferent sensory neurons, the disruption of dopaminergic and GABAergic signaling is the dominant mechanism associated with motor movement. The effects on important and broadly conserved signaling mechanisms in vertebrates suggest that NDL PCBs may contribute to neurodevelopmental abnormalities in humans and, with the startle response being critical for the survival of fish, to evolutionary adaptation in wildlife.

## Introduction

Humans and wildlife are exposed to a wide range of contaminants in the environment and the role of these chemical pollutants in developmental neurotoxicity, cognitive impairment, and neurodegenerative disorders has become a global environmental health concern (Landrigan et al. 2018). Among the top industrial-derived contaminants thought to be involved in developmental neurotoxicity are polychlorinated biphenyls (PCBs) (Boix et al. 2010; Boucher et al. 2009; Grandjean and Landrigan 2014). For decades, PCBs have been used as insulators, coatings, and caulks, and have persisted to date in the environment where they have globally accumulated in wildlife tissue and human cord blood, amniotic fluid, and mother’s milk (Ennaceur et al. 2008; Lancz et al. 2015a). PCBs are comprised of planar, dioxin-like (DL) congeners that lack ortho-chlorine substituents as well as non-planar, non-dioxin-like (NDL) congeners which have ortho-chlorine substituents. The levels of NDL PCBs in human and environmental samples are far greater than the levels of DL congeners, with NDL PCB153 and PCB138 being generally the most abundant. In human cord blood, PCB153 concentrations range from 3.7 to 200 ng g^-1^ lipid (Bergonzi et al. 2009; Herbstman et al. 2007; Lancz et al. 2015b; Patel et al. 2018).

During the early stages of life, the nervous system is generally more sensitive to chemical exposure compared to fully developed adults (Rice and Barone 2000), and therefore it is essential to identify the neurotoxic mechanisms that underpin the behavioral effects of environmental exposure to different PCBs during gestation and early life stages if we want to successfully treat and remediate health effects in humans and wildlife. Early developmental exposure to PCBs is associated with many neurodevelopmental disorders. For instance, increased serum PCB concentrations in children are associated with deficits in cochlear functions (Jusko et al. 2014; Trnovec et al. 2010), fine motor skills (Ribas-Fitó et al. 2001; Walkowiak et al. 2001), and cognitive impairment (Boucher et al. 2009; Chen et al. 1992). In rats, developmental exposure to NDL PCBs has been linked to behavioral changes, including hyperactivity, impulsiveness, and impairment of cognitive function (Berger et al. 2001; Boix et al. 2010; Johansen et al. 2014). NDL PCB developmental exposure also adversely affect the auditory function in adult rats, including elevated thresholds of brainstem auditory evoked potentials, auditory deficits at low frequencies, audiogenic seizures, and reduced auditory startle responses (Crofton et al. 2000; Goldey et al. 1995; Lilienthal et al. 2011, 2015; Poon et al. 2015). The observed effect on the auditory-evoked startle response is particularly relevant as it plays a critical role in the performance and survival of fish and amphibians, and involves cellular functions, sensory-motor pathways, and monoamine neurotransmitters that, ranging from fish to humans, are conserved among vertebrates (Stewart et al. 2014).

To date, mechanistic studies of neurodevelopmental effects of NDL PCBs suggest three major targets: (1) alterations in neurotransmission, particularly involving dopamine and serotonin, and consequently disruptions to Ca^2+^ signal transduction (Campagna et al. 2011; Dervola et al. 2015; Enayah et al. 2018; Fonnum and Mariussen 2009; Langeveld et al. 2012), (2) loss of outer hair cells (Crofton et al. 2000; Trnovec et al. 2010), and (3) altered axonal or dendritic morphogenesis (Keil et al. 2019; Lein et al. 2007; Pruitt et al. 1999; Yang et al. 2009). The links between these PCB targets and neurobehavioral deficits remain unknown. We used the zebrafish auditory system (**Figure 1**) as a model to decipher the involvement of these three major targets in disrupting the startle response after developmental exposure to NDL PCB153. Many of the pathways and genes that regulate startle behavior in fish also operate in human neurological disorders (Howe et al. 2013; Maximino and Herculano 2010; Meserve et al. 2020; Tropepe and Sive 2003; Wolman et al. 2015). We demonstrate that environmentally relevant concentrations (10 nM) of NDL PCB153 alter startle response which coincided with axonal swelling and dysregulated dopamine metabolite and GABA levels. Co-exposures to the dopamine D2-receptor antagonist and GABA receptor modulator drug haloperidol alleviated the delay in startle response. While similar responses were observed for other di-*ortho* NDL PCBs and the mono-*ortho* congener PCB118, a delayed startle response was not observed with the non-*ortho* PCB126, a potent AHR agonist. These results highlight the potential of low concentrations of NDL PCBs to disrupt broadly conserved neurotransmission tied to a mechanosensory deficit.

**Figure 1.**
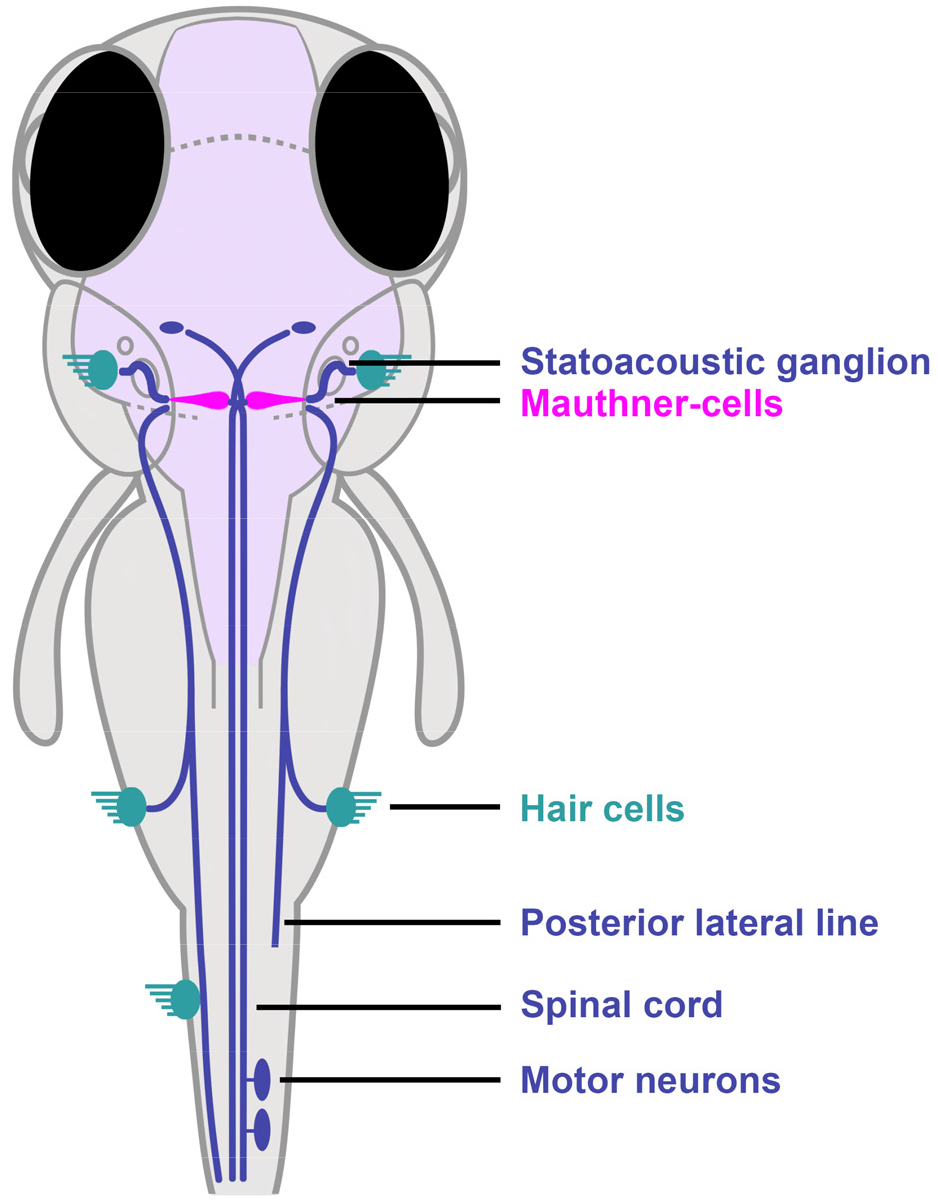
Startle response circuit in zebrafish. The proper execution of the auditory-evoked startle response relies on hair cells to sense the auditory stimulus, Mauthner-cells to integrate the signals, and motor neurons and musculature to execute the startle behavior.

## Results

### Morphological Effects

Exposure to contaminants during development can result in reduced growth, which can affect kinematics in swimming behavior (Voesenek et al. 2018). Zebrafish larvae exposed to 1000 nM PCB153 developed normally with no difference in body length at 6 dpf in comparison to the control group, in three separate trials (Trial 1: *t*(25.98) = .661, *p* = .514; Trial 2: *t*(37.98) = .443, *p* = .660; Trial 3: *t*(37.43) = .645, *p* = .523; **Figure S1, Supplemental Material**).

### Behavioral effects

Rohon-Beard cells are mechanosensory neurons that appear earliest during development in zebrafish. They are located in the dorsal region of the embryonic spinal cord and appear around 24 hpf (Roberts 2000). A few days into development, Rohon-Beard cells also relay sensory information to the M-cells (Liu and Hale 2017). In our experiments, a tactile stimulus on the trunk at 26 hpf evoked a C-shaped body flexion in both the DMSO- and the 1000 nM PCB153-treated embryos (*U* = 183, *p* = .599; **Figure S2**), indicating that the formation and function of Rohon-Beard cells were not impaired by PCB153.

Spontaneous and light-dark-stimulated swimming activity assessed at 6 dpf revealed that larvae exposed to PCB153 (1000 nM) had lower cumulative locomotion activity over ten minutes (*M* = 1182 mm, *SD* = 382) in comparison to the control group (*M* = 1510 mm, *SD* = 402), *t*(132.4) = 4.854, *p* < .0001 (**Figure S3A**). Zebrafish larvae typically increase locomotion in dark conditions following light exposure. Once stimulated by a rapid loss of illumination, both treatment groups exhibited similar swimming activity (**Figure S3B**).

The vibro-acoustic startle response includes two distinct motor behaviors: a short-latency C-bend (SLC) initiated within 5-15 ms of the stimulus, or a long-latency C-bend (LLC) beginning from 15-100 ms after the stimulus (Burgess and Granato 2007). The mean latency value calculated as the average of all SLC and LLC responses in each treatment population was 21 ± 20 ms for control larvae at the highest vibro-acoustic stimulus intensity (**Figure 2A, 2B**). By contrast, the NDL PCB-treated larvae that responded to the stimulus showed significantly longer average latencies in response to the same stimulus intensity given the controls, *H*(8) = 368.6, *p* < .0001. Larvae exposed to PCB153 responded at mean latencies that increased with dose. Thus at 10 nM PCB153 the latency was 26 ± 27 ms, at 30 nM it was 50 ± 36 ms, at 100 nM it was 50 ± 30 ms, and at 1000 nM the latency was 68 ± 32 ms. Results with the lowest PCB153 concentration tested (10 nM) were not significantly different from controls (*p* = .752). Larvae exposed to other NDL congeners also showed long latencies; with PCB138 it was 52 ± 37 ms, with PCB118 it was 45 ± 27 ms, and with PCB52 it was 39 ± 32 ms. Unlike the NDL PCB-treated larvae, larvae treated with the potent DL PCB126 had a mean latency of 22 ± 22 ms, essentially identical to that in the controls (*p* =.999).

**Figure 2.**
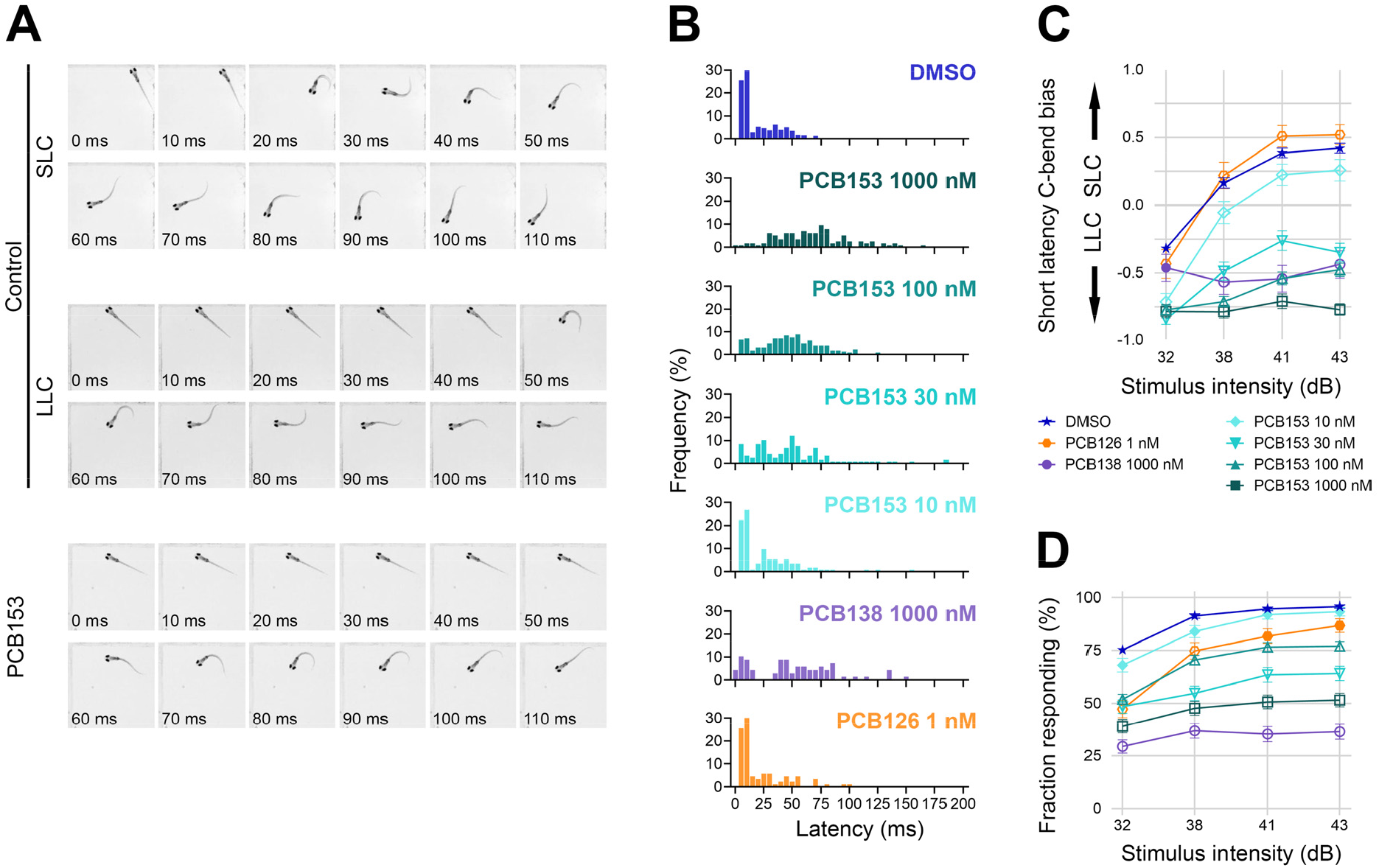
Effect of PCBs on startle response in 6 dpf zebrafish. (**A**) Representative short-latency C-bend (SLC) and long-latency C-bend (LLC) startle response of a control zebrafish larva at 6 dpf in comparison to a typical response of a PCB153 treated larva. (**B**) Vibro-acoustic startle latency at highest stimulus intensity (43 dB) in vehicle control (*n* = 479), NDL PCB153 1000 nM (*n* = 179), NDL PCB153 100 nM (*n* = 253), NDL PCB153 30 nM (*n* = 152), NDL PCB153 10 nM (*n* = 119), NDL PCB138 (*n* = 139), and DL PCB126 (n = 97) exposed larvae. **(C)** Bias of SLC and LLC and (**D**) response rate of different exposure groups at different vibro-acoustic intensities. Values are presented as mean ± SEM. All data points are biologically independent replicates from at least three independent experiments. Significant differences to DMSO controls (*p* < 0.01) are indicated by open symbols

Examining the SLC and the LLC responses separately can give insight into the underlying mechanism of effect. A greater stimulus intensity increases the probability of evoking an SLC, shifting the response bias from LLC to SLC (Troconis et al. 2017). Consistent with that we observed a shift to SLC with increasing stimulus intensity in solvent control (DMSO-treated) larvae (**Figure 2C, Figure S4A**). Larvae were tested 4 times at each stimulus intensity. A larva is categorized as having SLC bias if it responded with an SLC 3 or 4 times out of the 4 tests. At the highest stimulus intensity that we tested (43 dB), the majority of control larvae (300/448) responded with an SLC within 7 ± 3 ms. There were 43 larvae that showed no bias towards SLC or LLC (2 times SLC and 2 times LLC, bias = 0). There also were larvae that did not perform an SLC (*n* = 105) but did exhibit an LLC, with a mean latency of 54 ± 29 ms (**Figure 2C**). In contrast, all NDL PCB-treated larvae retained a response bias towards LLC at all stimulus intensities tested, which was significantly different from the control (*p* < .01 for the three highest stimulus intensities). However, as we decreased the PCB153 dose, there was an increased shift towards an SLC, yet even at 10 nM PCB153 the bias toward LLC was significantly different from the control (*p* <.01 for all stimulus intensities). PCB153 exposures starting at 24 hpf, after many structures of the brain and the neural tube have developed (Smith and Kimelman 2019), had the same effect on startle latency as exposures starting at 4 hpf (**Figure S4B)**.

The proportion of control larvae responding to the stimulus (with either an SLC or an LLC) increased from 75.2 ± 33.0% (*n* = 450) to 95.8 ± 17.0% (*n* = 457) from the lowest to highest stimulus intensity. In contrast, the proportion of PCB153-, PCB138-, and PCB52-treated larvae that responded remained below 70% at all intensities and were significantly different (*p* < .01) from the control, except for the larvae exposed to 10 nM PCB153 (**Figure 2D, Figure S5A**). In contrast to larvae exposed to the NDL PCBs, the larvae exposed to the DL PCB126 and NDL PCB118 had a response latency and rate similar to the control larvae.

The kinematics of the C-bend angle differ for the SLC and LLC startle responses, with SLCs exhibiting larger bend angles and faster angular velocities than LLCs (Burgess and Granato 2007). We examined the bend angles separately in SLC and LLC occurring after exposure to each PCB. Generally, there were increased bend angles associated with SLC response after NDL PCB exposure, even when there were very few SLC responders. Similarly, there were generally decreased bend angles in the LLC responders after NDL PCB exposures (**Figure S6**).

### Cellular structures

The proper execution of the startle response relies on hair cells that sense the auditory stimulus, M-cells to integrate the signals, and motor neurons and musculature to execute the startle behavior. Morphological analysis showed that M-cells, motor neurons, and musculature all appeared unaffected by PCB153, but the neurons innervating the hair cells appeared to be swollen (**Figure 3**). The L3 neuromast volume of control larvae was on average 569 µm^3^ (*SD* = 125), whereas larvae exposed to PCB153 (1000 nM) had significantly larger L3 neuromast volumes 848 µm^3^ (*SD* = 185), *t*(48.93) = 6.765, *p* < .0001 (**Figure 3B**). However, the number of hair cells in L3 neuromasts (**Figure 3C**) and the intensity of FM1-43 dye internalized by hair cells (**Figure 3D**) remained unchanged in PCB153-treated larvae in comparison to the control.

**Figure 3.**
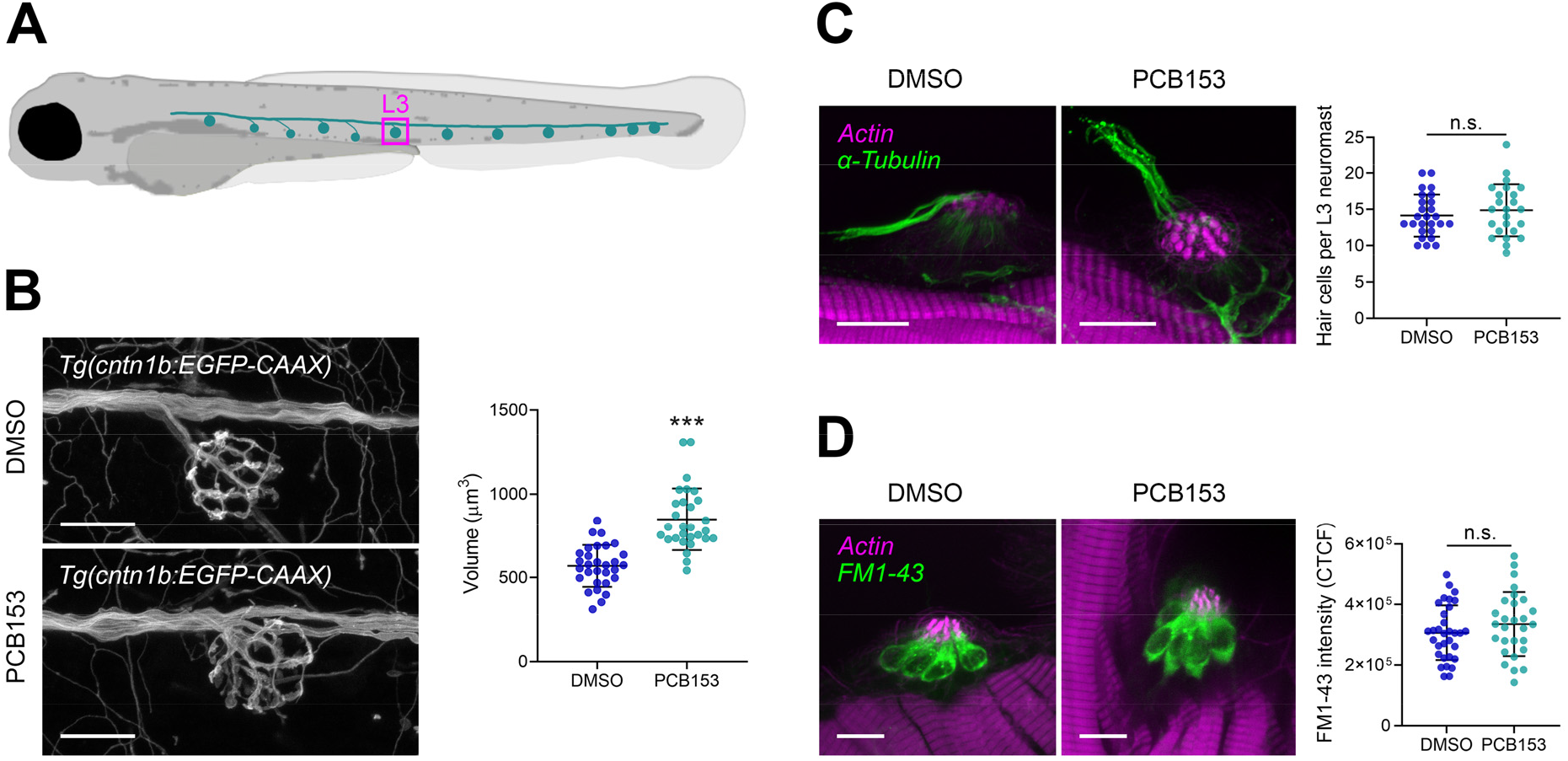
Effect of PCB153 on morphology and function of lateral line hair cells and innervating neurons. (**A**) Diagram indicating L3 neuromast. (**B**) Hair cell innervating neurons at neuromast L3 shown using *Tg(cntn1b:EGFP-CAAX)*. Volume of hair cell innervating neurons of DMSO (*n* = 30) and PCB153 (*n* = 29) treated larvae. Scale bar = 20 µm. (**C**) Whole-mount immune staining of kinocilia and hair cell innervating neurons (anti-acetylated tubulin antibody; green) and counterstain of stereocilia bundles and muscle tissue (actin; magenta) of DMSO (*n* = 26) and PCB153 (*n* = 24) treated larvae. The hair cell number was determined based on the number of kinocilia. Scale bar = 10 µm (**D**) Uptake of FM-143 dye as indication of functional hair cells of DMSO (*n* = 32) and PCB153 (*n* = 28) treated larvae. Scale bar = 10 µm. All data points are biologically independent replicates from three independent experiments. Asterisks indicate significant differences to controls (***p < 0.001).

The presence of functional M-cells is necessary for the SLC startle response. Staining with Mab 3A10 confirmed the presence of M-cells in all PCB153-exposed larvae (21/21) (**Figure 4A**). We tested the functional responsiveness of the M-cells by applying electric field pulses to head-restrained larvae. This electrical stimulation (**Figure 4B**) bypasses the sensory system and directly activates the M-cell (Tabor et al. 2014). There was no difference detected in response time between control and 1000 nM PCB153-exposed larvae (*U* = 907.5, *p* = .489), demonstrating that the signal transmission from the M-cell to the motor neurons is not disrupted in the PCB153-exposed fish (**Figure 4C**). The fraction responding also was similar in both the DMSO-treated (290/299) and the PCB153-treated (209/219) groups (GLM, *p* = .427; **Figure 4D**). However, the angle of the C-bend evoked by the electrical stimulation was larger in PCB153-exposed larvae (*M* = 101.4, *SD* = 38.60) than in the control larvae (*M* = 83.12, *SD* = 24.23). An unpaired t-test with Welch’s correction indicated that this difference was statistically significant, *t*(74.27) = 2.679, *p* = .0091 (**Figure 4E**). This difference could indicate a PCB153-evoked functional alteration of the M-cell axon or the motor neuron.

**Figure 4.**
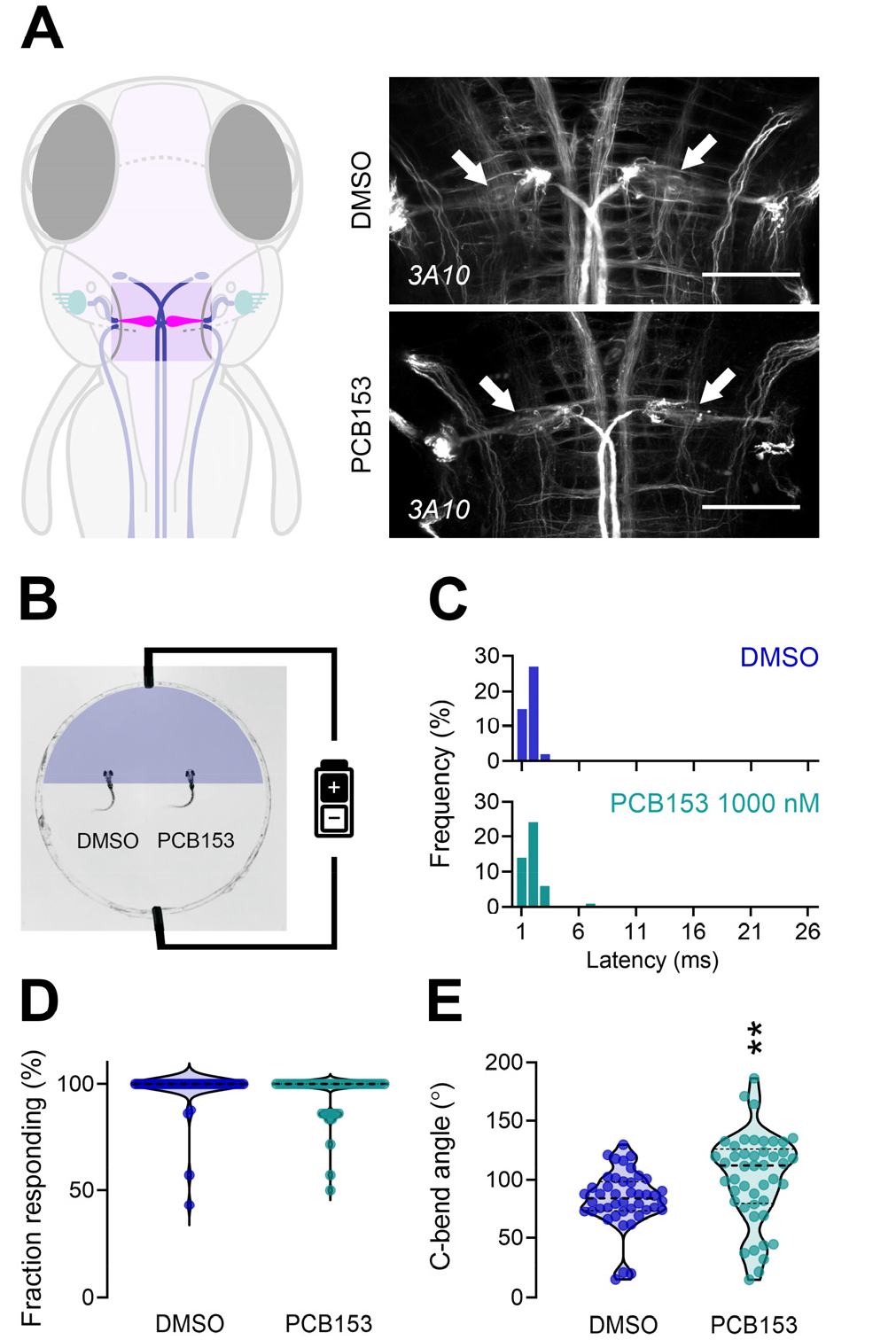
Effect of PCB153 on morphology and function of the Mauthner-cells. (**A**) Diagram indicating localization of Mauthner-cells in the hindbrain and brain tissue immunostaining with anti-neurofilament 3A10 antibody labeling reticulospinal neurons in control and PCB153 treated larvae at 6 dpf (ventral view). Mauthner (arrow) cells were present in all brain tissues (control: *n* = 23, PCB153: *n* = 21). Scale bar = 50 µm. (**B**) Diagram of electrical stimulation performed to assess the functionality of Mauthner-cells. Both DMSO (*n* = 44) and PCB153 (*n* = 45) treated larvae show similar (**C**) latency to electrical stimulation and **(D)** response frequency (seven stimuli per larvae), indicating that the M-cells are present and functional. (**E**) The increased C-bend angle in PCB153 exposed larvae may be indicative of impaired motoneuron or muscle development. All data points are biologically independent replicates (one value representing the mean of seven consecutive electric field pulses) from three independent experiments. Dashed lines represent median and quartiles and asterisks indicate significant differences to controls (**p < 0.01).

For rapid and effective axonal signal transmission, proper myelination of axons is key. We did not find PCB153 to reduce cells of the oligodendrocyte lineage in the spinal cord of the *Tg(olig2:EGFP)* transgenic line treated with PCB153, (*t*(36) = .553, *p* < .584). We also did not find any differences in myelination itself in the *Tg(mbp:EGFP-CAAX)* transgenic line treated with PCB153 (**Figure S7**).

Proper function and activation of the motor neurons are crucial for the completion of the startle circuit, leading to muscle contractions and the characteristic swimming behaviors. The main caudal primary (CaP), middle primary (MiP), and rostral primary (RoP) motor neurons did not show any distinct structural differences between exposed and control larvae (**Figure S8**). The fast muscle fibers in the myotome were not differentially striated in the PCB153 treatment group (**Figure S9**). No structural difference in the motor neurons or the myotome was apparent in PCB153-exposed larvae, suggesting that the increased bend angle does not originate from any overt structural differences in the motor neurons or muscle fibers.

### Neuronal activity

The release of neurotransmitters into the synaptic cleft is a characteristic process in signal transmission. Both serotonergic and dopaminergic neuromodulators can affect startle latency (Jain et al. 2018), and evidence has been found that NDL PCBs alter these neuromodulators in mammals (e.g. Castoldi et al. 2006; Enayah et al. 2018; Wigestrand et al. 2013). We observed that PCB153 (1000 nM) altered the levels of several neurotransmitters in 6 dpf zebrafish larvae (**Figure 5A**). Significantly increased levels of GABA (*t*(6) = 9.719, *p* < .001), choline-chloride (*t*(6) = 8.459, *p* < .001), 3-methoxytyramine hydrochloride (*t*(6) = 3.803, *p* = .009), and L-valine (*t*(6) = 3.134, *p* = .020) were measured in PCB-exposed larvae in comparison to control larvae; while L-Glutamine levels were decreased in comparison to control larvae (*t*(6) = 5.966, *p* < .001).

**Figure 5.**
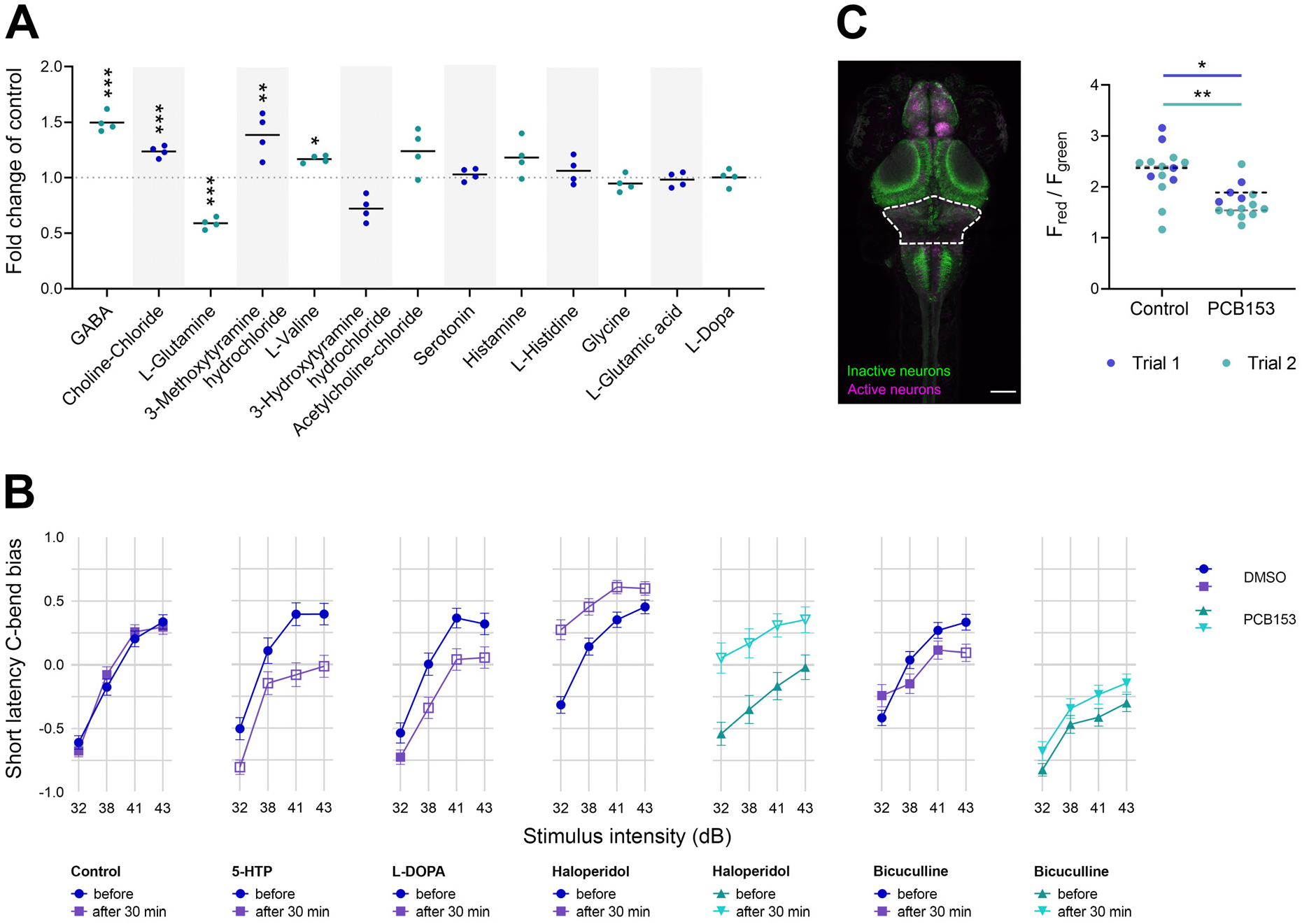
Involvement of neurotransmitters in startle latency. (**A**) Neurotransmitter levels in 6 dpf larvae exposed to 1000 nM PCB153 in comparison to control (*n* = 4). (**B**) Startle bias shift induced by 30-minute exposure to serotonin, dopamine, and GABA neurotransmitter modulators in 6 dpf control and 100 nM PCB153 exposed larvae. Control (*n* = 138-172), 5-HTP (*n* = 67-80), L-DOPA (*n* = 76-94), haloperidol (Control *n* = 123-162, PCB153 *n* = 37-61), and bicuculline (Control *n* = 90-162, PCB153 *n* = 83-137). All data points are biologically independent replicates from at least three independent experiments. Significant differences to respective controls (*p* < 0.01) are indicated by open symbols. (**C**) Representative maximum intensity z-projection from confocal stack after photoconversions of freely swimming 6 dpf CaMPARI larvae. Scale bar = 100 µm. Ratio of red to green fluorescence intensity in the rostral hindbrain (indicated by dashed line in image) in DMSO (*n* = 14) and 1000 nM PCB153 (*n* = 14) exposed startling larvae in 4 °C (cold) medium. Dashed lines indicate median per trial. Two independent experiments. Asterisks indicate significant differences to controls (*p < 0.05, **p < 0.01, and ***p < 0.001).

To assess the interaction between PCB exposure and changes in neuromodulatory activity, we pharmacologically manipulated the neurotransmitter levels using dopamine, serotonin, and GABA receptor agonists and antagonists. In control larvae, a 30 min exposure to dopamine precursor L-DOPA and D2-receptor agonist Quinpirole, as well as serotonin precursor 5-HTP, shifted the response to LLC, at multiple intensities tested (*p* < .01; **Figure 5B**). In contrast, the dopamine antagonist haloperidol exposure induced a shift to SLC in controls across all stimulus intensities (*p* < .01; **Figure 5B**). Haloperidol induced the same shift to SLC in PCB153-exposed larvae (*p* < .001), suggesting that the startle circuit is functional but repressed by PCB (**Figure 5B**). The response rate remained unchanged after exposure to neurotransmitter modulators except for 5-HT, which caused an increase in response probability across all stimulus intensities in control larvae (*p* < .01); quinpirole, which caused a decrease in response probability at the two lowest stimulus intensities in control larvae (*p* < .01); and haloperidol, which caused an increase in response probability at 41 dB in PCB153-treated larvae (*p* < .01; **Figure S5**). It is noteworthy, that co-exposure to L-DOPA and PCB153 throughout development increased the startle response rate of PCB153-treated larvae significantly (*p* < .01; **Figure S5C**), yet only induced a significant shift towards SLC at 38 dB (*p* < .01; **Figure S4C**).

In the fish brain, a distinct population of serotonergic neurons is found in the posterior tuberculum/hypothalamus, forming a prominent horseshoe-like pattern that is one of several brain regions where 5-HT is synthesized (Oikonomou et al. 2019). To assess whether the development of serotonergic cells in the brain of PCB153 larvae was affected, we performed immunohistochemistry using an antibody for 5-HT (**Figure S10**). The intensity of the fluorescent signal in the hypothalamus was not different, comparing PCB153 (*M* = 4.70 x 10^6^, *SD* = 2.10 x 10^6^) with the control (*M* = 4.38 x 10^6^, *SD* = 8.1 x 10^5^) group. This suggests that the serotonergic neurons in this region were not affected.

During specific behaviors or sensory experiences, only a small percentage of neurons are active. We analyzed the active neuronal cell populations in PCB153-treated and -untreated larvae, focusing on the rostral hindbrain where the M-cells are located, to assess whether there were changes in neural activity. We examined neuronal activity using the CaMPARI zebrafish line, which contains a protein that upon UV illumination permanently photoconverts from green to red fluorescence in cells with high calcium levels (Fosque et al. 2015). This genomic tool enabled us to look for active neuronal cell populations in larvae during normal swimming movement and in larvae experiencing a vibroacoustic-stimulated startle response. At room temperature (24 °C) no difference in the red to green ratio (F_red_/F_green_) was measured comparing control and PCB153-treated larvae, neither when swimming nor startling (**Figure S11**). However, when stimulating high neuronal activity by placing the larvae in cold (4 °C) medium, swimming control larvae exhibited a significantly higher F_red_/F_green_ ratio (*M* = 2.30, *SD* = 0.51) in comparison to swimming PCB153-treated larvae (*M* = 1.70, *SD* = 0.31) on average across two trials (**Figure 5C**). Importantly, the difference between control and PCB153-treated larvae swimming in cold medium was significant in both trials (trial 1: *F*(1, 18) = 121, *adj. p* = .028 and trial 2 *F*(1, 32) = 65.2, *adj. p* < .001). Similar differences in neuronal activity between control and PCB153 treated larvae were measured for startling larvae (**Figure S11**).

## Discussion

While its use has mostly ended decades ago, PCBs remain a globally important and persistent group of pollutants that continues to impact both wildlife and humans. Prenatal exposure to PCBs is now recognized to contribute to neurobehavioral disorders in humans and potentially other organisms, yet the mechanisms that underlie its neurotoxicity remain largely unknown (Klocke and Lein 2020). Using developmental exposure of zebrafish to PCB153, we identify here an imbalance in dopamine metabolites and GABA mediating mechanosensory deficits in fish, encompassing hearing and touch. The behavioral effect was found to be specific to NDL PCBs and at concentrations relevant for both humans and aquatic life. These findings highlight a so far unrecognized impact of commonly observed concentrations of NDL PCBs on the escape response in fish and link a conserved neurotoxic mechanism to behavioral deficits.

We observed a delayed startle response for all NDL PCBs tested and subsequently examined a number of morphological features of the underlying startle circuit anatomy (hair cell, afferent neurons, M-cell, oligodendrocyte lineage cells, myelination, motor neuron, muscle; **Figure 1**), as potentially affected by the NDL PCB153. Only the volume of the neurons that innervate the hair cells showed a measurable deviation from the control condition in PCB153-exposed larvae (**Figure 3**). We thus infer that the enlargement of axon termini is contributing to the neurotoxic potential of NDL PCBs. This is an understudied target of PCB toxicity with only a few studies associating NDL PCB exposure with altered dendritic arborization (Klocke and Lein 2020; Lein et al. 2007; Yang et al. 2009). The direct links to neurobehavioral changes remain to be shown. However, swelling of afferent terminals is associated with hearing deficits in both mammalian models (Puel et al. 1991) and zebrafish (Sebe et al. 2017). Axonal swelling is also a pathology appearing in many neurodegenerative diseases (Coleman 2005), thus exposure to NDL PCBs may be a contributing factor in such diseases.

The proper function of the M-cell is essential in transmitting the signal that leads to the rapid startle response. At the structural level, laser-ablation of the M-cell abolishes SLC responses without affecting LLC responses (Lacoste et al. 2015). Our finding that NDL PCB-exposed larvae predominantly exhibit LLC without loss of M-cells (**Figure 2C, 4A**), suggests that the M-cells are receiving a delayed input, are not firing properly, or that the signal propagation downstream of the M-cell is disrupted. To distinguish these, we directly activated the M-cells by applying electrical stimulation to larvae with their heads restrained and found that, unlike auditory stimulation, PCB153-exposed fish had latencies indistinguishable from the controls (**Figure 4C**). Furthermore, direct stimulation of PCB153-exposed larvae resulted in a normal response rate (**Figure 4D**), although the bend angle was significantly increased (**Figure 4E**). These results indicate a proper function of M-cells in NDL PCB-treated larvae. However, the changes in bend angles we observed during these responses could indicate alterations downstream of the M-cells.

There is little known about the mechanisms leading to an aberrant turning angle. Since we found no obvious structural impairment downstream of the M-cell (oligodendrocyte lineage cells, myelination, motor neuron, musculature), we speculate that the increased bend angle in electrically stimulated larvae exposed to PCB153 may be linked to a sub-cellular disruption such as gap junctions. It is known that PCBs can alter gap junction function in cells of many organs (Bager et al. 1997; Kang et al. 1996; Machala et al. 2003) including NDL PCB specific inhibition of gap junctional intercellular communication in neural crest cells (Nyffeler et al. 2018) and neuronal stem cells (Kang et al. 2001). Gap junctions are intercellular channels formed by connexins (Cx). RNA sequencing of zebrafish larvae exposed to PCB153 in a similar exposure regime (4-120 hpf) and concentration (10 µM) significantly altered the expression of connexin genes cx28.9, cx42, and cx35b (Aluru et al. 2020). Cx35 is expressed on the Mauthner neuron lateral dendrites in zebrafish (Jain et al. 2018) and plays a crucial role in regulating spinal motor activity during fast motor behavior of adult mosquitofish (Serrano-Velez et al. 2014). We thus propose that gap junction function could be disrupted, which may be a future study direction using single-cell RNA sequencing or genetically encoded optogenetics that has yet to be developed in a model organism (Dong et al. 2018).

Neurotransmitters also play a key role in the startle response. A previous screening of more than 1000 pharmacologically active compounds identified serotonergic and dopaminergic modulators among the largest class of drugs that shift bias between SLC and LLC startle behavior (Jain et al. 2018). Specifically, acute exposure to the D3R agonist (7OHD) shifts the startle bias from LLC to SLC, while exposure to a D3R antagonist (or a 5-HT1AR agonist) shifts bias toward LLC behavior (Jain et al. 2018). Our results extend these findings, showing that 30 minutes of exposure to L-DOPA or 5-HTP induced a shift from SLC to LLC at higher stimulus intensities, as opposed to the D2R antagonist haloperidol, which evoked a shift to a stronger SLC bias at all stimulus intensities (**Figure 5B**). NDL PCBs are thought to affect dopaminergic pathways (Mariussen and Fonnum 2001; Tanaka et al. 2018; Wigestrand et al. 2013), which is partly supported by our measurements of catecholamines and traces amines. Specifically, we found that the extracellular dopamine metabolite 3-methoxytyramine hydrochloride and the inhibitory transmitter GABA were significantly increased (**Figure 5A**). The interplay between dopamine and GABA receptor activity is complex. In the mammalian system, dopamine is a direct-acting modulator of GABA_A_Rs (Hoerbelt et al. 2015). The simultaneous increase of a dopamine metabolite and GABA in PCB153-exposed zebrafish larvae may indicate that increased levels of GABA are a downstream effect of dysregulated dopamine activity. With both dopamine and GABA playing critical roles in motor activity, we hypothesize that PCB153 impairs dopaminergic neurotransmission and subsequently increases GABA levels, leading to the predominant LLC response.

Furthermore, we propose that GABA is involved in the swelling of the afferent terminals (**Figure 3**) as GABA activates Cl^-^ channels leading to an influx of Cl^-^ accompanied by H_2_O (Cesetti et al. 2012; Rungta et al. 2015). The authors report that the osmotic neuronal swelling alters neuronal activity. This study thus provides new evidence that GABA is involved in the neurobehavioral mode of action of higher chlorinated NDL PCBs, as has been shown for lower chlorinated congeners (Fernandes et al. 2010).

An important finding of this study is that PCB153-exposed larvae can execute SLC when treated with haloperidol, a D2-antagonist that can increase dopamine turnover in acute exposure (Magnusson et al. 1987), suggesting dopamine turnover as the major disrupted mechanism causing the sensory deficit. There is a large body of evidence showing that NDL PCBs, including PCB153, modulate dopaminergic neurotransmission by inhibiting the dopamine precursor enzyme tyrosine hydroxylase and dopamine synthesis itself, in both neural crest cells and adult laboratory animal models including non-human primates (Dervola et al. 2015; Enayah et al. 2018; Fonnum and Mariussen 2009; Seegal et al. 1990). In zebrafish larvae, PCB153 exposure during development elicited differential expression of nine genes related to the dopamine pathway, including tyrosine hydroxylase 2 (th2) (Aluru et al. 2020). Yet linking such effects to behavior is rare. Tanaka et al. (2018) suggested that the NDL PCB dominant mixture Aroclor 1254 and BDE-47 functionally inhibited dopaminergic neurons in early zebrafish embryos, as increased activity in 26 hpf embryos was inhibited by supplementation with L-tyrosine and L-DOPA (Tanaka et al. 2018).

The restoration of the delayed startle phenotype by haloperidol, together with the increased levels of a dopamine metabolite and GABA in PCB153-exposed larvae (**Figure 5A**), indicates that disruption of dopaminergic and GABAergic signaling between neurons of the auditory circuit is the dominant effect of PCB153-mediated aberrant motor behavior. These signaling processes are widely conserved across vertebrate species and are important in human neurological diseases. For example, a recent study found that multiple gene mutations causing defective startle response in zebrafish were also associated with disorders of the locomotor system in humans (Meserve et al. 2020).

The localization of the PCB-induced disruption of dopaminergic signal transmission is likely downstream of the hair cells, potentially in the hindbrain. Although dopaminergic neurons innervate lateral line neuromasts, where D1R antagonist reduces the hair cell activity (Toro et al. 2015), we found no difference in the hair cell activity of PCB153-exposed larvae (**Figure 3D**). This is in contrast to the hindbrain, where neuronal activity was reduced in PCB153 exposed CaMPARI zebrafish larvae (**Figure 5C**), indicating a reduced capacity for release or turnover of calcium. Intracellular Ca^2+^ signaling in neurons controls neurotransmitter secretion among many other cellular processes and thus may be indirectly involved in dopamine or GABA regulation. Some NDL PCBs (PCB 95, 138) are known to inhibit voltage-gated calcium channels (Langeveld et al. 2012) contributing to interference with calcium homeostasis and consequently neurotransmission and synaptic plasticity (Brunelli et al. 2012; Campagna et al. 2011; Gafni et al. 2003; Ta et al. 2006).

The startle response in fish and amphibians is a fundamental mechanism involved in predator escape. PCB153-exposed zebrafish larvae predominantly responded with an LLC, which not only delays the response, increasing the chance to be caught by a predator but also usually does not displace the animal from its original location (**Figure 2A** and Jain et al., 2018). While survival of predator attack is likely context-dependent and species-specific, quick escape responses increase chances of survival (McCormick et al. 2018), suggesting the highly abundant di-*ortho*-NDL PCBs may provide a substantial selection pressure and thereby could drive evolutionary adaptation. We have observed a significant shift to LLC at concentrations of PCB153 as low as 10 nM (3.6 µg L^-1^). PCBs are associated with sediments, where concentrations of PCB153 frequently reach multiple µg kg^-1^ (Lai et al. 2015). PCB concentrations in organisms may be even higher due to extensive bioconcentration; killifish from New Bedford Harbor have 6.5 to 7 µg of PCB153 per g wet weight (Gräns et al. 2015). All the *ortho*-NDL PCBs tested in this study increased the startle latency. Thus, in the environment, where PCBs occur in mixtures of numerous congeners that likely have additive effects, the combined effect could occur at concentrations lower than that of any single congener. Taken together, these results highlight the importance of the neurotoxic potential of NDL PCBs in the natural environment.

Based on the comprehensive approach presented in this study, it can be inferred that PCB153 may alter dopamine signaling, leading to an increase in GABA that ultimately results in swelling of axonal terminals in the auditory circuit. The dysregulation of neurotransmitters is likely the dominating mechanism for the behavioral deficit, as the treatment with the dopamine antagonist haloperidol restored the startle capability of PCB153-treated fish. While the involvement of connexins remains speculative, our findings provide a foothold into the dopamine dysregulating potential of PCB153 and likely other NDL PCBs in sensory-motor processing in developing vertebrates, and link that to motor behavior. In a direct comparison, the present study suggests that the adverse behavioral effects observed are induced by all NDL PCBs tested, but not by the classical DL PCB126. Whether this is true for compounds that are structurally similar to NDL PCBs, such as some PBDEs, is not known and will require further research. A key finding is that the effect of NDL PCBs on the startle latency occurs at concentrations relevant for both environmental and human exposures, thereby exposing a mechanism that is relevant to both a behavior with inherent selection pressures in fish and neurotoxicological diseases in humans. This study therewith provides valuable clues for tackling health effects in humans and wildlife caused by environmental exposure to widespread PCBs.

## Materials and Methods

### Animal model rationale

Zebrafish have relatively simple neural circuits, particularly for the auditory system. Many of the components of the auditory circuit are homologous in mammals and fish, including hair cells, neural transmission, and lower motorneurons, while the mechanosensory lateral line system and Mauthner (M) cells are exclusively found in fish and a cochlea is only found in mammals. In fish, vibro-acoustic stimuli are first sensed by the hair cell bundles in the auditory, vestibular, and lateral-line systems and are rapidly encoded into trains of action potentials in the afferent neurons (**Figure 1**) (Hudspeth 1985). The signal detected through the sensory hair cells of the lateral line is transmitted through the anterior and posterior lateral line ganglia whereas the auditory input is transmitted through the VIIIth statoacoustic nerve, all of which synapse to the lateral dendrite of the M-cell and two segmental homologs (Nakayama 2004; Pujol-Martí and López-Schier 2013). The signal is propagated down the M-cell axon in the spinal cord to activate the motoneurons leading to a unilateral contraction of the trunk musculature.

### Zebrafish lines and husbandry

Zebrafish (*Danio rerio*) were maintained at 28.5 °C and on a 12:12 h light-dark cycle according to the guidelines of the Zebrafish Model Organism Database (ZFIN, http://zfin.org) and institutional guidelines at WHOI and the experimental procedures approved by the Institutional Animal Care and Use Committee (ID Number BI21981.01). Adult zebrafish were fed brine shrimp and Gemma Micro 300 pellets (Skretting) every day. In this study, we used wild-type zebrafish of the AB strain obtained from Zebrafish International Resource Center (ZIRC), as well as several transgenic lines: the photoconvertible calcium indicator line *Tg[elavl3:CaMPARI(W391F+V398L)]*^*jf9*^ kindly provided by Jessica Plavicki (Brown University, USA), and the transgenic lines *Tg(cntn1b:EGFP-CAAX)* (Panlilio et al. 2021) labeling axons, *Tg(olig2:EGFP*)^*vu12*^ (Park et al. 2007; Shin et al. 2003) marking oligodendrocyte precursor cell bodies, *Tg(sox10:mRFP)* (Takada et al. 2010) marking oligodendrocyte lineage cells, as well as *Tg(mbp:EGFP-CAAX)* (Almeida et al. 2011) *and Tg(mbp:EGFP)* (gift from Dr. Kelly Monk, generated by Dr. Charles Kaufman in the laboratory of Dr. Leonard Zon, Harvard Medical School, Boston, MA) marking myelin sheaths. Zebrafish eggs were obtained by pairwise breeding and kept in 0.3x Danieau’s (17 mM NaCl, 0.2 mM KCl, 0.12 mM MgSO_4_, 0.18 mM Ca(NO_3_)_2_ and 1.5 mM HEPES, pH 7.6) in glass vials at a density of 10 individuals per 10 mL, at a constant water temperature of 28 ± 1°C and standard 14-h light/10-h dark cycle. The medium was supplemented with 1-Phenyl-2-thiourea (PTU, 0.003% w/v) from 24 hours post-fertilization (hpf) onwards to prevent pigment formation when necessary for imaging.

### Early-life stage exposure

Fertilized eggs were used for exposures starting at 4 hpf to solvent control (0.1% DMSO, v/v), 2,2′,4,4′,5,5′-hexachlorobiphenyl (PCB153), 2,2′,5,5′-PCB, 2,2′,5,5′-tetrachlorobiphenyl (PCB52), 2,3’,4,4’,5-pentachlorobiphenyl (PCB118), 2,2’,3,4,4’,5’-hexachlorobiphenyl (PCB138), or 3,3’,4,4’,5-hexachlorobiphenyl (PCB126). Additionally, one exposure with PCB153 was started at 24 hpf, to assess whether effects are induced during or after initial neuronal cell differentiation; sensory neurons send their projection along the ectoderm as early as 16 hpf (Kimmel 1993). The lowest exposure concentration for PCB153 was 10 nM (3.6 ng mL^-1^), estimated to result in ∼5 ng g^-1^ (5 ppm) per zebrafish larva, based on comparison to PCB95 uptake levels (Ranasinghe et al. 2019). PCB153 concentrations found in one study of maternal cord blood serum were 0.27 ng mL^-1^ or 110 ng g^-1^ lipid (Lancz et al. 2015b) while other reported concentrations range from 3.7 to 200 ng g^-1^ lipid (Bergonzi et al. 2009; Herbstman et al. 2007; Lancz et al. 2015b; Patel et al. 2018). We also used higher exposure concentrations of PCB153 including 30 nM, 100 nM, and 1000 nM, which are found in fish of contaminated ecosystems such as the New Bedford Harbor in Massachusetts, USA (Gräns et al. 2015). For PCB52, PCB118, and PCB138, a concentration of 1000 nM was chosen, to compare to the highest effect concentration of PCB153. The DL-PCB126 is cardioteratogenic in the low nanomolar range. To avoid malformations, a concentration of 1 nM was chosen, which is still highly effective at activating the aryl hydrocarbon receptor and the expression of AHR target genes. Exposures were refreshed daily until four days post-fertilization (dpf) and all larvae were kept in Danieau’s from 5 to 6 dpf. All imaging and behavioral assays were performed at 6 dpf, except for the touch assay, which was performed at 26 hpf. For all behavioral assays, only larvae with an inflated swim bladder were used.

The influence of neurotransmitters on the startle behavior was assessed by exposures to compounds known to interfere with monoamines. At 6 dpf, larvae were evaluated for their startle behavior and then immersed in the dopamine precursors L-DOPA (1 mM) or L-tyrosine (1 mM), dopamine receptor agonist quinpirole (10 µM), dopamine D2-receptor antagonist and GABA receptor modulator haloperidol (10 µM), endogenous serotonin 5-HT (10 µM), and serotonin precursor 5-HTP (1 mM) for 30 min in the well plate before assessing the startle behavior again. Additionally, the effect of neurotransmitter modulation during development on startle response was assessed by exposing larvae from 3 to 120 hpf to L-DOPA (1 mM), L-tyrosine (1 mM), and quinpirole (10 µM).

### Behavioral assays

Larval locomotion was monitored using the Noldus Daniovison system (Noldus Information Technology, Leesburg, VA, USA) using 6-dpf larvae distributed in 48-well plates (1 larva per well). After an acclimation period of 40 minutes in the illuminated chamber, spontaneous swimming activity was tracked for 10 minutes. Following the spontaneous swimming, three alternating cycles of dark and light conditions (10 minutes each) were run. The experiment was repeated three times with cohorts from separate breeding events (*n* = 24 per group and breeding event). Video data were recorded with 30 frames per second via an infrared camera. Obtained data were analyzed with the supplied software EthoVision XT® 12 (Noldus. Inc.).

The activity of Rohon-Beard sensory neurons was tested at 26 hpf by lightly touching DMSO- and 1000 nM PCB153-exposed embryos (*n* = 10 per group and breeding event) using a piece of fishing line affixed to a glass pipette. Each embryo was touched ten times and the response was scored as follows: 0, no response; 0.5, light tail flicking response, 1, coiling in the opposite direction of the touch stimulus, yielding a score of a maximum of 10 per embryo. The experiment was repeated twice with cohorts from separate breeding events.

Vibro-acoustic startle response was assessed as described previously (Panlilio et al. 2021). Briefly, the acoustic stimuli were delivered by a vibrational exciter producing acoustic vibrations at a frequency of 1000 Hz for a duration of 2 ms at four different amplitudes (32, 38, 41, 43 dB). For each amplitude, the stimulus was delivered four times, spaced 20 seconds apart, to a 4 x 4 well-plate mounted on top of the speaker, each well containing an individual larva. The response was tracked with a high-speed camera (Edgertronic SC1) at 1000 frames s^-1^ for 250 ms (13 ms before and 237 ms after the stimulus). PulsePal was used to synchronize the camera recording to the acoustic stimuli using a TTL pulse. Automated analysis of larval movement kinematics (response frequency, latency, turning angle) was performed using the FLOTE software package (Burgess and Granato 2007). Each treatment group was assessed at least three times with eggs from different clutches. The total replicate numbers are stated in **Table S1 (Supplemental Material)**.

A startle response in fish can be executed with a short latency C-bend (SLC) or long latency C-bend (LLC), with the SLC being the more frequent response at higher stimulus intensities. The cut-off time between SLC and LLC can vary depending on environmental conditions such as temperature and has been determined empirically for this experiment using a Gaussian mixture model (Panlilio et al. 2020; **Figure S12**). Responses within 15 ms were categorized as SLC, latencies greater than 15 ms as LLC. Following this classification, the median response type (Relative Startle Bias) for each responsive fish was determined for each stimulus intensity (32, 38, 41, and 43 dB). Bias per individual fish was calculated as (frequency of SLC – frequency of LLC) / total responses. Pure SLC responses result in a value of +1, while pure LLC responses are a value of -1 (Jain et al. 2018).

Electric field pulses (EFPs) directly activate the M-cells (Tabor et al. 2014). M-cell functionality was assessed by using head-restrained larvae (*n* = 46 per treatment group). For each run, one control and one PCB-exposed larva had their heads embedded in 1.5% agarose, side-by-side, with their tails free to move. Seven consecutive EFPs (4.4 V cm^-1^ for 2 ms, square pulse) were applied with a recovery time of 20 s in between pulses. Response frequency, latency, and bend angle were tracked using FLOTE.

### Larval growth

Larval length (from snout to the tail-fin end) was measured in DMSO (*n* = 57) and 1000 nM PCB153 (*n* = 51) exposed larvae using ImageJ. The experiment was performed three times with embryos from different clutches.

### FM1-43 labeling of functional hair cells

The vital fluorescent dye FM1-43 (n-(3-triethylammoniumpropyl)-4-(4-(dibutylamino)-styryl) pyridinium dibromide; Invitrogen) specifically labels hair cells in the lateral line and inner ear of vertebrates by entering through open mechanotransduction channels and filling the cytoplasm of hair cells (Meyers et al. 2003). At 6 dpf, zebrafish larvae of 1000 nM PCB153-exposed (*n* = 32) and unexposed (*n* = 28) groups were immersed in Danieau’s supplemented with 3 µM FM1-43 for 30 s, rinsed three times in Danieau’s, anesthetized in 1 mM 3-amino-benzoic acid (Tricaine), and mounted in 1.5% low melting agarose for confocal imaging of the L3 neuromast. The experiment was performed three times independently.

### Visualization of M-cells, kinocilia, and muscle fibers, neurons, oligodendrocyte lineage cells, and neuronal activity

Whole-mount immunohistochemistry of zebrafish larvae and brains was performed following standard procedures. Briefly, 10 AB larvae per exposure group were treated as described above and subsequently fixed in 4% paraformaldehyde (PFA) in PBS overnight at 4 °C. For labeling brain tissue, larval brains were dissected before continuing with the protocol. Multiple rinsing steps in PBS containing either 0.5% Triton-X100 or 0.1% Tween-20 (PBST) for larvae or brains, respectively, was followed by an antigen retrieval step in 150 mM Tris HCl (pH 9.0) at 70 °C (Inoue and Wittbrodt 2011), and a permeabilization step using proteinase K (10 µg/mL for 3 minutes). Subsequently, the samples were blocked in blocking solution (10% Normal Goat Serum, 1% DMSO, 1% BSA, 88% PBST) for 1 h and then stained with blocking solution supplemented with the primary antibody for three days at 4 °C. The primary anti-neurofilament mouse monoclonal antibody 3A10 (Developmental Hybridoma Bank; 1:100 dilution) was used to label a subset of hindbrain spinal cord projecting neurons such as the M-cell neurons (DMSO *n* = 23, PCB153 *n* = 21). The anti-5HT rabbit polyclonal antibody (Sigma-Aldrich, #S5545, 1:500) was used to label serotonergic cells (DMSO *n* = 16, PCB153 *n* = 13). The anti-acetylated alpha-tubulin mouse polyclonal antibody (Sigma-Aldrich, #T6793, 1:1000 dilution) was used to label kinocilia and hair cell innervating neurons (DMSO *n* = 25, PCB153 *n* = 24). After several washing steps, in PBST, the samples were incubated with Alexa Fluor 488- or 594-conjugated goat-anti-mouse or goat-anti-rabbit secondary antibodies (Abcam, 1:500 dilution) overnight at 4 °C and rinsed several times in PBST before mounting in ProLong Diamond Antifade mountant (Invitrogen) for imaging. For visualization of skeletal muscle fibers (DMSO *n* = 20, PCB153 *n* = 20), larvae were fixed as above, followed by permeabilization in 5% Triton-X for overnight and incubation in phalloidin Alexa-546 conjugate (Invitrogen, 1:500). All staining experiments were repeated three times with larvae from independent clutches.

The *Tg(cntn1b:EGFP-CAAX)* transgenic line was used to image the cell bodies of neurons that innervate the hair cells at the L3 position in the lateral line (DMSO *n* = 22, PCB153 *n* = 20, two independent experiments). The axons of the main caudal primary motoneurons were visualized using the *Tg(olig2:EGFP)* transgenic line (DMSO: *n* = 21, PCB153: *n* = 18). The double transgenic *Tg(olig2:EGFP) x Tg(sox10:mRFP)* was used to quantify oligodendrocyte lineage cells in the spinal cord of in DMSO (*n* = 22) and 1000 nM PCB153 (*n* = 16) treated larvae (Shin et al. 2003). All larvae were anesthetized in tricaine (0.16% MS222) before mounting laterally in 1.5% low melting agarose on a glass-bottom microscopy dish (MatTek). Fluorescently labeled cell bodies were imaged at the distal end of the intestine, around the L3 neuromast. All experiments were performed three times independently.

Neuronal activity in DMSO- and PCB153-treated larvae was visualized using the CaMPARI (*elavl3*:CaMPARI) transgenic zebrafish line with an engineered fluorescent protein that permanently photoconverts from green to red in the presence of elevated calcium and UV light (Fosque et al. 2015). Photoconversion was achieved by mounting an LED light (405 nm; LEDSupply) between the speaker and a custom-made 3-well plate containing the freely swimming larvae (DMSO- and PCB153-treated). Larvae of the same treatment were divided into two batches: one was photoconverted in medium at room temperature (RT) and the other one at 4°C to elicit high neuronal activity. Photoconversion was achieved by shining the LED light for 10 sec either with one vibrational stimulus of 43 dB at the start or without stimulus. Subsequently, larvae were anesthetized, embedded in agarose, and whole brains imaged using the 10x objective. The experiment was performed twice (*n* = 11-16 per treatment group).

### Image acquisition and processing

Whole-brain, hair cells, M-cell, and motor neurons were imaged on a Zeiss LSM 800 confocal system with a 10x/0.45 Plan-Apochromat, 20x/0.8 Plan-Apochromat, or 63x/1.2 C-Apochromat water immersion objective. Airy scan was applied for hair cell imaging at 63x. Z-stacks were merged to a single plane by maximal intensity projection using Fiji (Schindelin et al. 2012).

The number of *olig2* cells dorsal of the spinal cord (DMSO *n* = 22, 1000 nM PCB153 *n* = 16) was counted in confocal image stacks using the 3D projection of the Zeiss Zen blue software. Myelination was evaluated after imaging the spinal cord (DMSO *n* = 26, 1000 nM PCB153 *n* = 27) using the 20x objective and scored for deficits according to Panlilio et al. 2020. Representative myelination was imaged at 40x. The intensity of serotonergic neurons in the brain (DMSO *n* = 16, 1000 nM PCB153 *n* = 13) was measured performing a Sum Slices Z-projection followed by drawing a ROI around the 5-HT immuno-responsive cells, and per image, three random squares (area size = 10.65 x 10.65) within the non-fluorescent area of the ROI for background subtraction (for orientation see **Figure S12**). Similarly, the FM1-43 intensity was measured drawing a ROI around the fluorescent hair cells and measuring the background in three random squares (DMSO *n* = 20, 1000 nM PCB153 *n* = 20). Corrected total cell fluorescence (CTCF) was calculated according to the formula: Integrated density - (ROI x Mean fluorescence of background readings). For quantification of active neurons in the brain of freely swimming and startling CaMPARI zebrafish larvae, Sum Slices Z-projection were analyzed for red fluorescence intensity in the rostral hindbrain (for orientation see **Figure S13**). To calculate the volume of hair cell innervating neurons, automatic image thresholding with fixed minimum and maximum threshold values (30; 255) using the Otsu method) was applied to all images. Subsequently, image stacks were analyzed using an imageJ macro looping through the stack and summing the area measurements, then multiplying this sum by the depth of each slice (DMSO *n* = 30, 1000 nM PCB153 *n* = 29). The presence of primary motor neurons was confirmed using the Simple Neurite Tracer plugin in Fiji (DMSO *n* = 21, 1000 nM PCB153 *n* = 18). All image evaluations and quantifications were done without the knowledge of the experimental condition of the specimens.

### Neurotransmitter analysis

The analysis of neurotransmitters, including histamine, L-histidine, L-glutamic acid, gamma-aminobutyric acid (GABA), acetylcholine, L-DOPA, 3-methoxytyramine, serotonin, 5-hydroxyindole-3-acetic acid, glycine, L-glutamine, L-valine, 3-hydroxytyramine and choline, was performed with a method developed in a previous study (Zhao et al. 2020). Briefly, the zebrafish embryo samples were spiked with 100 ppb isotope-labeled internal standards and then homogenized. The homogenized tissue samples were extracted with 200 μL of 80:20 v/v ACN/Milli-Q water containing 0.1% formic acid and 1% ascorbic acid and 200 μL ACN. After protein precipitation, the supernatants were dried and redissolved in 100 μL of Milli-Q water containing 0.1% formic acid for UHPLC-MS/MS analysis. Neurotransmitters were quantified by a high-performance liquid chromatography system coupled to a Q-Exactive plus quadrupole-Orbitrap mass spectrometer (UHPLC-Q-Orbitrap, Thermo fisher, US). The separation of neurotransmitters was carried out with an Acquity UPLC HSS T3 column with Milli-Q water containing 0.1% formic acid and ACN containing 0.1% formic acid as mobile phase. The MS acquisition was performed in the full scan mode and parallel reaction monitoring mode. The experiment was performed once (*n* = 4 per treatment group).

### Statistical Analyses

All statistical analyses were conducted using Graph Pad Prism 8.4 except some of the startle behavior analysis, which was completed in R. All quantitative data were tested for normality using the Shapiro-Wilk test, and if there were an unequal number of subjects between groups, the data were tested for homogeneity in variances using the Brown-Forsythe test. All treatment groups were compared to respective control groups.

Responsiveness to auditory stimuli (response or not) and startle type (SLC vs. LLC) was modeled in R (version 4.0.2) using a binomial generalized linear mixed effect model (glmer() from the lme4 R package) (Bates et al. 2015). The response to electrical stimulus was modeled using glm() from the lme4 R package. For the auditory response, we included treatment as a fixed effect and the replicate experimental trials as a random effect. The model specification for “fraction responding” was as follows: Responders ∼ TreatmentGroup + (TreatmentGroup|trial). The model specification for “startle bias” was as follows: SLCorLLCresponse ∼ TreatmentGroup + stimdB + (1|trial). The data were analyzed fitting the model for each stimulus intensity independently. The model specification for the electrical stimulation is as follows: Responders ∼ TreatmentGroup.

To assess treatment differences of normal data with three or more groups (fluorescence intensity CaMPARI), ANOVA comparisons were performed followed by a multiple comparison using the Tukey’s correction to adjust the critical values. A Tukey-corrected *p*-value < .05 was considered statistically significant. For normal data with heterogeneity within-group variances (LLC bend angle), a Welch ANOVA followed by a Dunnett’s T3 multiple comparison test was performed. For non-normal data with three or more groups (SLC bend angle), a Kruskal-Wallis followed by the Dunn’s multiple comparison test was performed. For normal data with two groups (length, mean latency, bend angle in e-stim, oligodendrocyte lineage cell count, 5-HT intensity), an unpaired two-tailed t-test with Welch’s correction (Welch’s t-test) was performed. The nonparametric scoring data of the touch assay and latency in e-stim was analyzed using a two-tailed Mann-Whitney test. The data are presented as mean ± *SD* with single data points (replicates) superimposed on the graph unless specified in the figure legend.

### CRediT authorship contribution statement

**Nadja R. Brun:** Conceptualization, Methodology, Formal analysis, Investigation, Writing - original draft, Writing - review & editing, Visualization, Funding acquisition. **Jennifer M. Panlilio:** Conceptualization, Methodology, Software, Formal analysis, Investigation, Writing - review & editing, Funding acquisition. **Kun Zhang:** Methodology, Investigation, Writing - review & editing, Funding acquisition. **Yanbin Zhao:** Methodology, Investigation, Writing - review & editing, Funding acquisition. **Evgeny Ivashkin:** Methodology, Investigation. **John J. Stegeman:** Conceptualization, Writing - review & editing, Funding acquisition. **Jared V. Goldstone:** Conceptualization, Writing - review & editing, Project administration.

## Acknowledgments

The authors would like to thank Neel Aluru for the helpful discussion and providing the confocal microscope and Ed Levin and Mark Hahn for critically reading the manuscript and making valuable suggestions.

