## Supplemental Material for "Neurological mechanism of sensory deficits after developmental exposure to non-dioxin-like polychlorinated biphenyls (PCBs)"

**Table S1.** Replicate number per treatment group used in the startle response assay and statistical results.

| **Treatment** | ***n*** | **In comparison to** | ***n*** | ***p* value for Fraction responding** | | | | ***p* value for SLC bias** | | | |
|---|---|---|---|---|---|---|---|---|---|---|---|
| | | | | 32 dB | 38dB | 41dB | 42 dB | 32 dB | 38dB | 41dB | 42 dB |
| DMSO | 479 [a] | 1000 nM PCB153 [b] | 139 | <.001 | <.001 | <.001 | <.001 | <.001 | <.001 | <.001 | <.001 |
| | | 100 nM PCB153 [b] | 234 | <.001 | <.001 | <.001 | <.001 | <.001 | <.001 | <.001 | <.001 |
| | | 30 nM PCB153 [b] | 147 | <.01 | <.001 | <.001 | <.001 | <.001 | <.001 | <.001 | <.001 |
| | | 10 nM PCB153 [b] | 103 | .59 | .603 | .787 | .147 | <.01 | <.01 | <.01 | <.01 |
| | | 1000 nM PCB138 [b] | 110 | <.001 | <.001 | <.001 | <.001 | 0.229 | <.001 | <.001 | <.001 |
| | | 1000 nM PCB118 [b] | 100 | .086 | <.001 | .223 | .025 | <.001 | <.001 | <.001 | <.001 |
| | | 1000 nM PCB52 [b] | 105 | <.001 | <.001 | <.001 | <.001 | .490 | <.01 | <.001 | <.001 |
| | | 1 nM PCB126 [b] | 83 | <.001 | <.01 | .751 | .088 | <.01 | <.001 | <.001 | <.001 |
| | | DMSO + L-DOPA 4-120 hpf [b] | 62 | .247 | .182 | .203 | .756 | .874 | .024 | <.01 | <.01 |
| | | DMSO + Quinpirole 4-120 hpf [b] | 125 | .562 | .827 | .61 | .395 | .330 | .158 | .877 | .289 |
| DMSO 24-120 hpf | 80 | PCB153 24-120 hpf | 83 | <.001 | <.001 | <.001 | <.001 | 0.248 | <.01 | <.001 | <.001 |
| DMSO + L-DOPA 4-120 hpf | 62 | PCB153 + L-DOPA 4-120 hpf | 139 | <.01 | <.001 | <.001 | <.001 | .349 | <.01 | .212 | .271 |
| DMSO 0 min | 161 | DMSO 30 min | 161 | .112 | .113 | .543 | .518 | .024 | .028 | .159 | .984 |
| PCB153 0 min | 58 | PCB153 30 min | 58 | .962 | .763 | .478 | .034 | . 723 | .114 | .894 | .960 |
| DMSO before 5-HT | 49 | after 30 min 5-HT | 49 | <.001 | <.001 | <.001 | <.001 | .329 | .863 | .986 | >.999 |
| DMSO before 5-HTP | 78 | after 30 min 5-HTP | 78 | .294 | .834 | .392 | .755 | <.001 | <.01 | <.001 | <.001 |
| DMSO before L-DOPA | 90 | after 30 min L-DOPA | 90 | .061 | .221 | .373 | .244 | .123 | <.001 | <.001 | <.001 |
| DMSO before L-tyrosine | 85 | after 30 min L-tyrosine | 85 | .635 | .539 | .32 | .64 | .239 | .155 | .111 | .171 |
| DMSO before Quinpirole | 101 | after 30 min Quinpirole | 101 | <.001 | <.001 | .169 | .020 | .122 | .671 | <.001 | <.01 |
| DMSO before Haloperidol | 177 | after 30 min Haloperidol | 177 | .617 | .457 | .104 | .86 | <.001 | <.001 | <.001 | <.01 |
| PCB153 before Haloperidol | 94 | after 30 min Haloperidol | 94 | .952 | .507 | <.001 | .978 | <.001 | <.001 | <.001 | <.001 |
| DMSO before Bicuculline | 163 | after 30 min Bicuculline | 163 | .144 | .636 | .303 | .156 | . 063 | .091 | .019 | <.001 |
| PCB153 before Bicuculline | 143 | after 30 min Bicuculline | 143 | .843 | .914 | .864 | .664 | .167 | .086 | .081 | .211 |
| DMSO before Nipecotic Acid | 152 | after 30 min Nipecotic Acid | 152 | .317 | .657 | .721 | .385 | .197 | .857 | <.01 | .260 |
| PCB153 before Nipecotic Acid | 166 | after 30 min Nipecotic Acid | 166 | .022 | .445 | .591 | .664 | .988 | .574 | .363 | .527 |

[a] total *n* for DMSO
[b] in comparison to DMSO of respective trial date

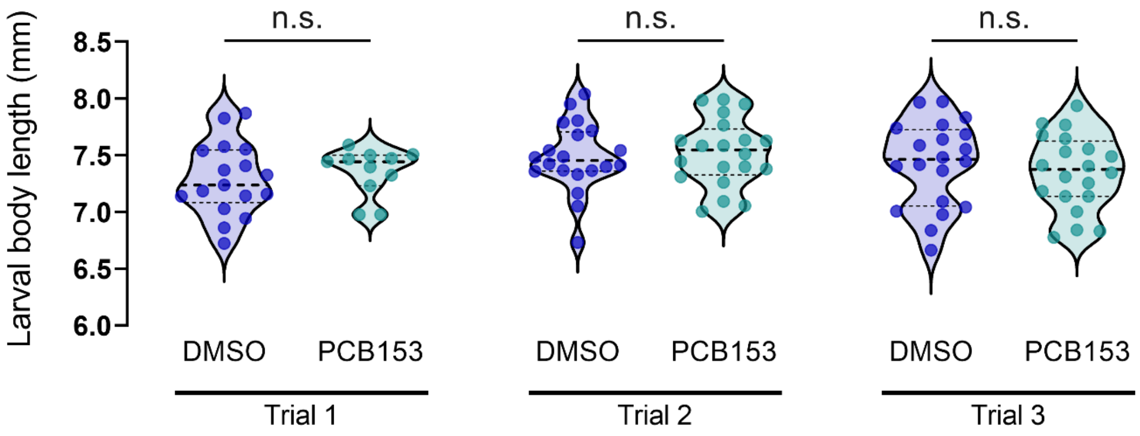


**Figure S1. Larval length.** No difference in larval length at six dpf was determined comparing solvent control with PCB153 treated larvae. Larval body length was measured from snout to tail-fin end. All data points are biologically independent replicates from three independent experiments (trials). Trial 1: DMSO $n$ = 17, PCB153 $n$ = 11, Trial 2: DMSO $n$ = 20, PCB153 $n$ = 20, Trial 3: DMSO $n$ = 20, PCB153 $n$ = 20. Dashed lines represent median and quartiles.

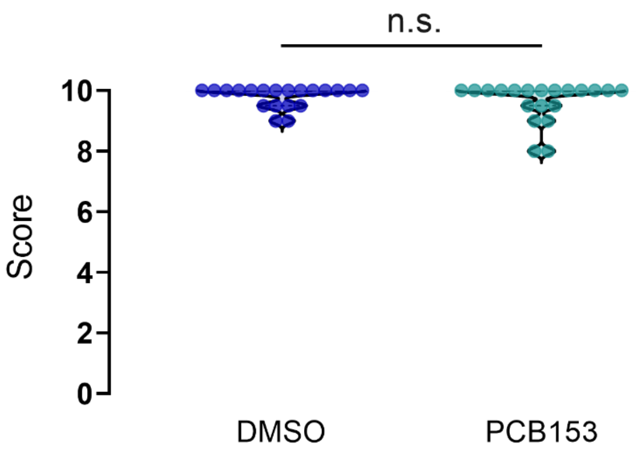


**Figure S2. Embryonic mechanosensory response.** The activity of Rohon-Beard cells in the trunk zebrafish embryos was tested by a light touch at 26 hpf. Each embryo was touched ten times and individual responses were scored as follows: 0, no response; 0.5, light response, 1, coiling in the opposite direction of the touch stimulus. Total score was calculated as the sum of individual scores. Statistical analysis revealed no difference between DMSO and 1000 nM PCB153 treated embryos (n = 20 per group). Data points represent biologically independent replicates from two independent experiments.

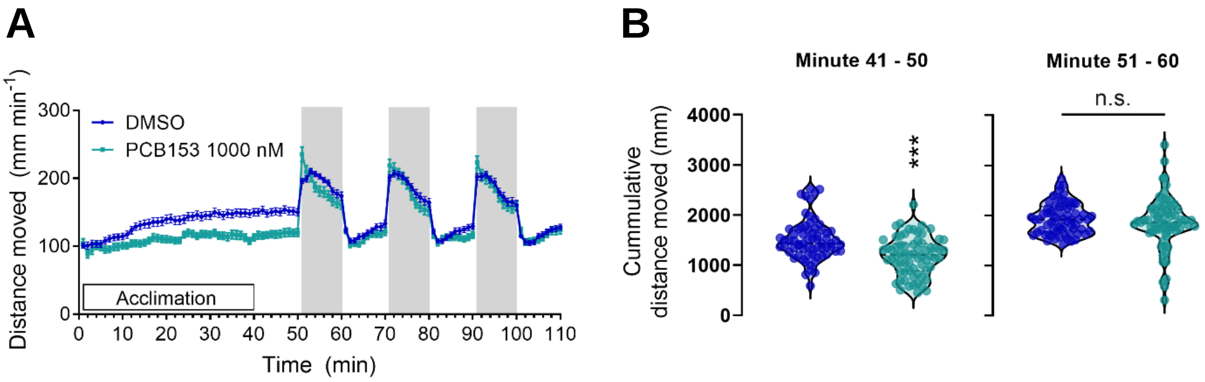


**Figure S3. Larval locomotion. (A)** Larval locomotion was tracked at six dpf over 120 minutes with three dark phases of 10 minutes each. Error bars represent the mean ± SEM. **(B)** The cumulative spontaneous swimming activity (41-50 min) of PCB153 ($n$ = 67) treated larvae is lower in comparison to the solvent control ($n$ = 68) treated larvae. No difference between control and treatment is observed in the alternating light/dark phases. All data points represent biologically independent replicates from three independent experiments.

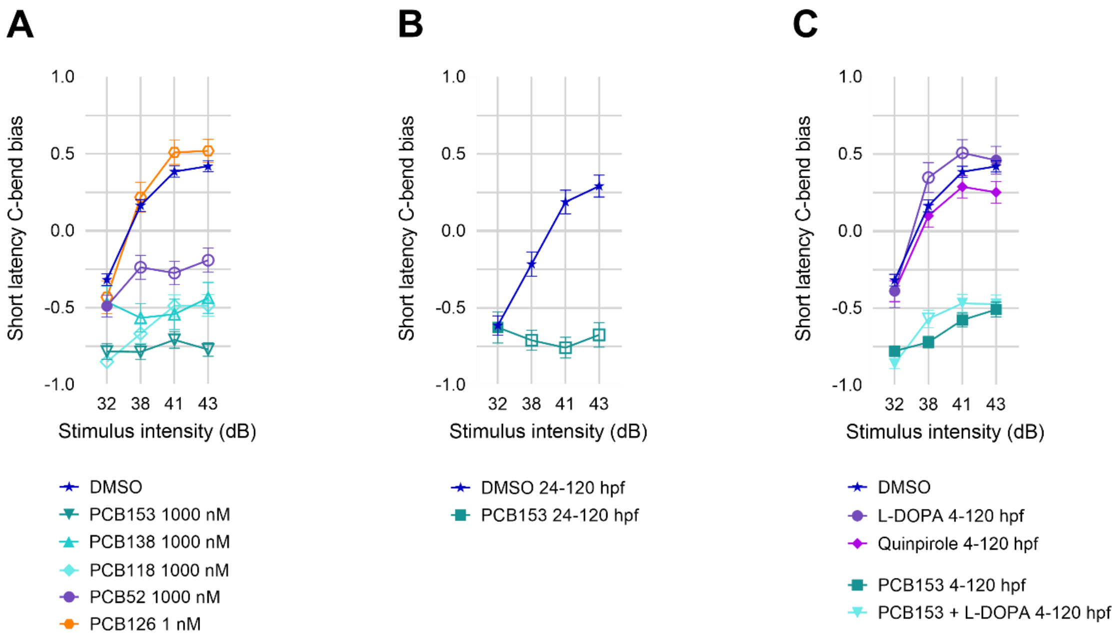

D

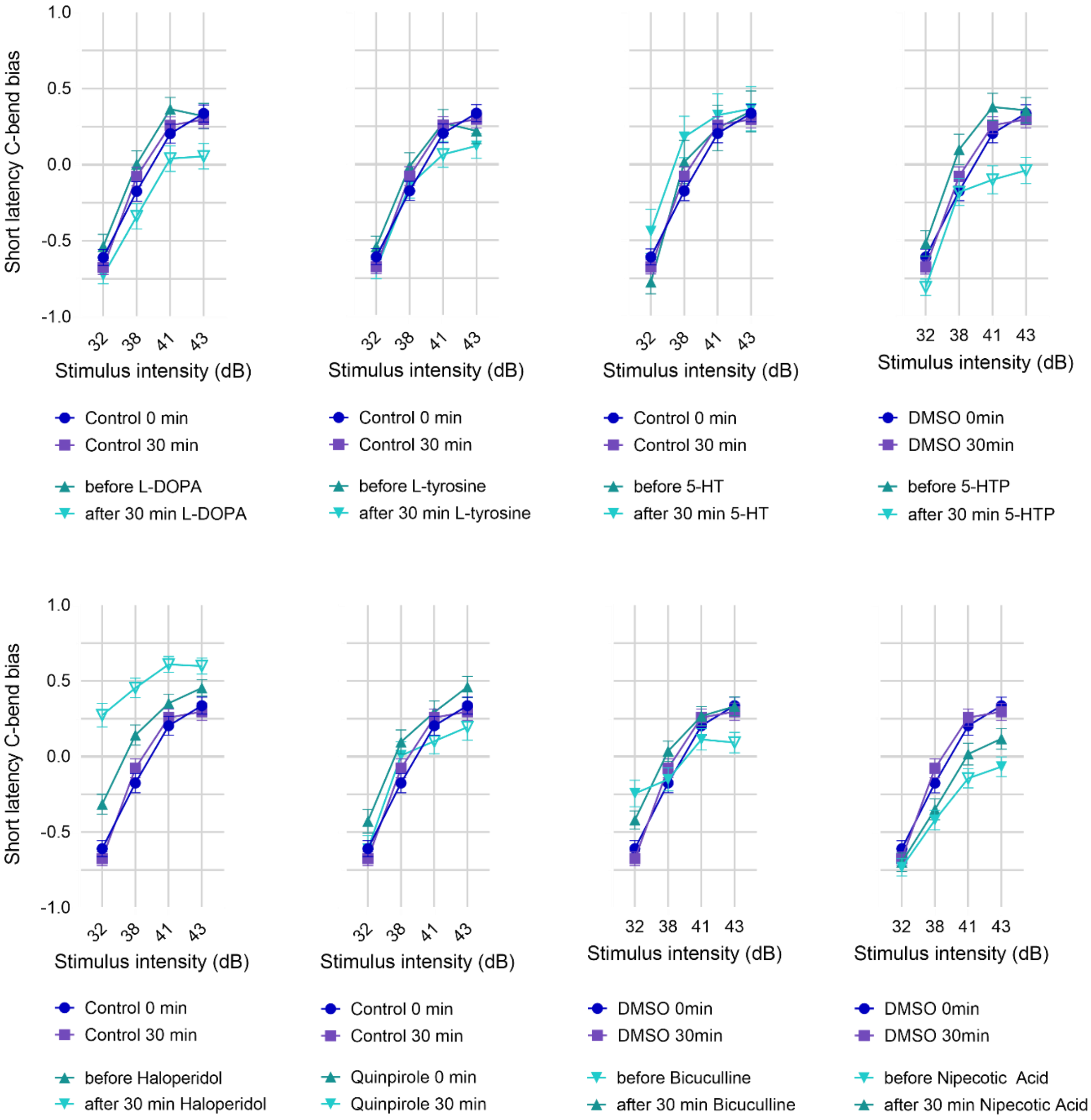

Short latency C-bend bias
1.0
0.5
0.0
-0.5
-1.0
32
38
41
43
Stimulus intensity (dB)
Control 0 min
Control 30 min
before L-DOPA
after 30 min L-DOPA
Control 0 min
Control 30 min
before L-tyrosine
after 30 min L-tyrosine
Control 0 min
Control 30 min
before 5-HT
after 30 min 5-HT
DMSO 0min
DMSO 30min
before 5-HTP
after 30 min 5-HTP
Control 0 min
Control 30 min
before Haloperidol
after 30 min Haloperidol
Control 0 min
Control 30 min
Quinpirole 0 min
Quinpirole 30 min
DMSO 0min
DMSO 30min
before Bicuculline
after 30 min Bicuculline
DMSO 0min
DMSO 30min
before Nipecotic Acid
after 30 min Nipecotic Acid


E

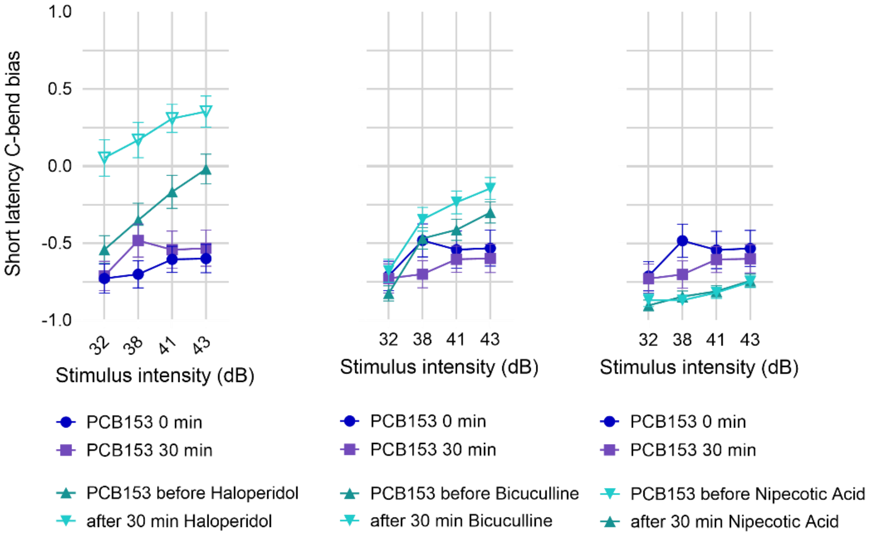

Short latency C-bend bias
1.0
0.5
0.0
-0.5
-1.0
32
38
41
43
Stimulus intensity (dB)
PCB153 0 min
PCB153 30 min
PCB153 before Haloperidol
after 30 min Haloperidol
PCB153 0 min
PCB153 30 min
PCB153 before Bicuculline
after 30 min Bicuculline
PCB153 0 min
PCB153 30 min
PCB153 before Nipecotic Acid
after 30 min Nipecotic Acid

**Figure S4. Short latency C-bend bias at different stimulus intensities.** Shifts from long latency C-bend (LLC) to short latency C-bend (SLC) with increasing stimulus intensities in control groups (DMSO *n* = 411-448, DMSO 24-120 hpf *n* = 73-80, DMSO 0 and 30 min *n* =138-172) in comparison to (**A**) different non-dioxin-like-PCBs (PCB153 *n* = 91-115, PCB138 *n* = 37-71, PCB118 *n* = 93-101, PCB52 *n* = 87-102) and the dioxin-like PCB126 (*n* = 63-86), (**B**) exposure to 1000 nM PCB153 excluding gastrulation (24-120 hpf; PCB153 *n* = 50-60), (**C**) exposure throughout development (4-120 hpf) to L-DOPA 1 mM (*n* = 51-66) or Quinpirole 10 µM (*n* = 109-122) and PCB153 100 nM alone (*n* = 198-228) and with co-exposure to L-DOPA (*n* = 117-142), (**D**) neurotransmitter modulators (L-DOPA *n* = 76-94, L-tyrosine *n* = 73-92, 5-HT *n* = 25-32, 5-HTP *n* = 67-80, Haloperidol *n* = 129-178, Quinpirole *n* = 63-109, Bicuculline *n* = 90-162, Nipecotic Acid *n* = 105-157 ) for 30 min in DMSO treated larvae at 6 dpf. (**E**) neurotransmitter modulators (Haloperidol *n* = 57-82, Bicuculline *n* = 83-137, Nipecotic Acid *n* = 83-156) for 30 min in 100 nM PCB153 treated larvae at 6 dpf and 100 nM PCB153 only (*n* = 37-47). Values are presented as mean ± SEM. All data points are biologically independent replicates from at least three independent experiments, *n* varies between stimulus intensities due to varying successful tracking numbers per video. Hollow symbols indicate a significant difference to respective controls ($p < 0.01$).

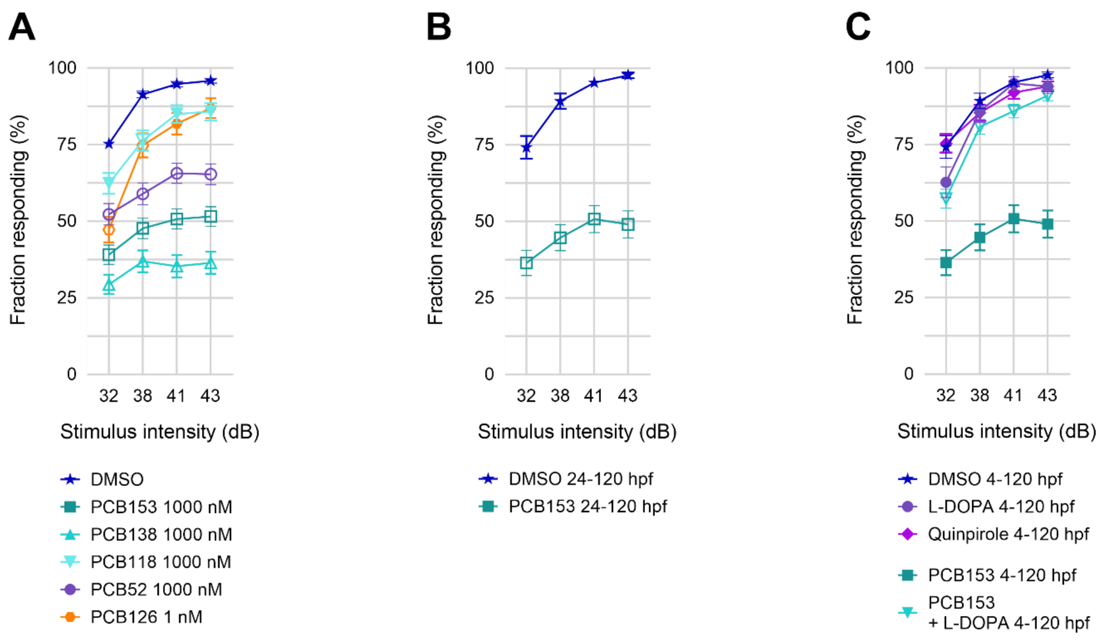

D

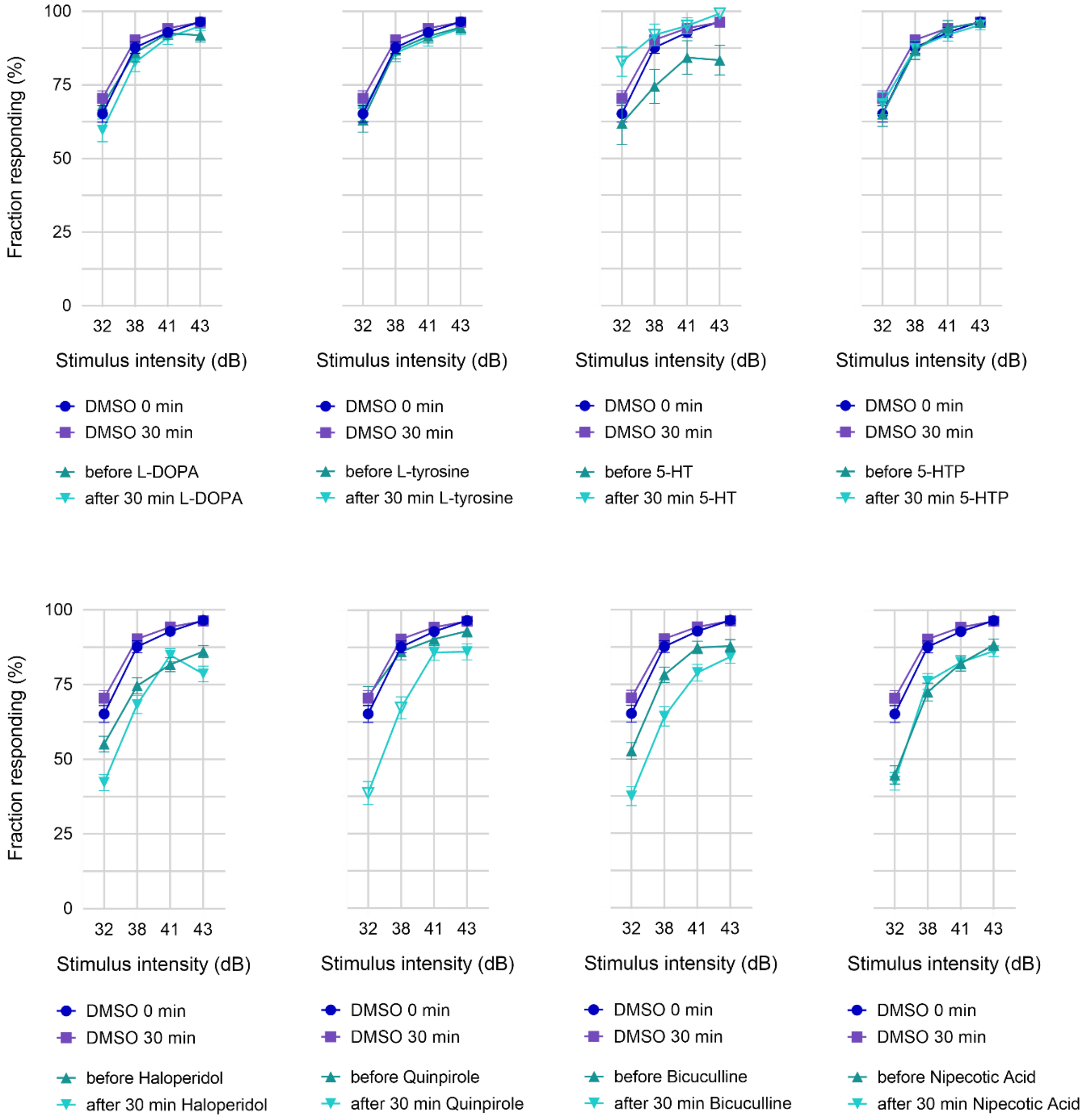

Fraction responding (%)
100
75
50
25
0
32 38 41 43
Stimulus intensity (dB)
DMSO 0 min
DMSO 30 min
before L-DOPA
after 30 min L-DOPA
before L-tyrosine
after 30 min L-tyrosine
before 5-HT
after 30 min 5-HT
before 5-HTP
after 30 min 5-HTP
before Haloperidol
after 30 min Haloperidol
before Quinpirole
after 30 min Quinpirole
before Bicuculline
after 30 min Bicuculline
before Nipecotic Acid
after 30 min Nipecotic Acid


E

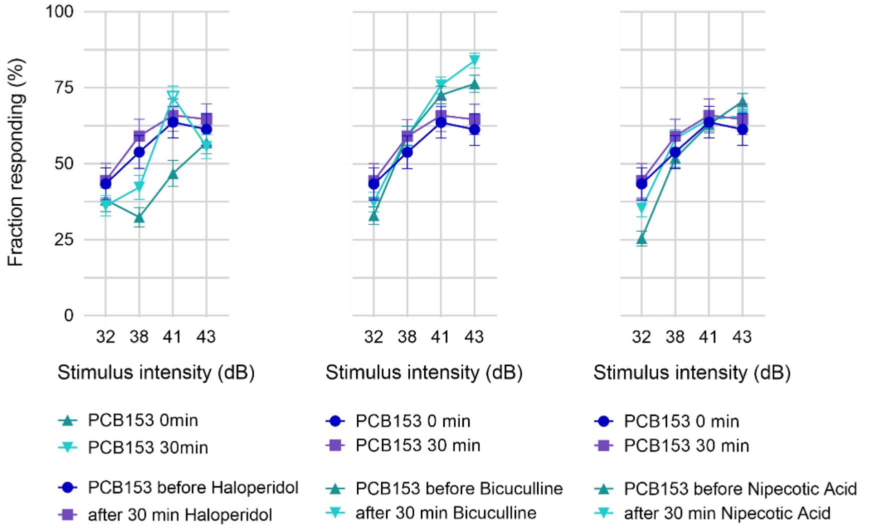

Fraction responding (%)
100
75
50
25
0
32 38 41 43
Stimulus intensity (dB)
PCB153 0min
PCB153 30min
PCB153 before Haloperidol
after 30 min Haloperidol
PCB153 0 min
PCB153 30 min
PCB153 before Bicuculline
after 30 min Bicuculline
PCB153 before Nipecotic Acid
after 30 min Nipecotic Acid

**Figure S5. Startle response rate at different stimulus intensities.** Response rate in control groups (DMSO *n* = 449-457, DMSO 24-120 hpf *n* = 79-80, DMSO 0 and 30 min *n* =158-175) in comparison to (**A**) different non-dioxin-like-PCBs (PCB153 *n* = 159-164, PCB138 *n* = 128-131, PCB118 *n* = 107-108, PCB52 *n* = 112) and the dioxin-like PCB126 (*n* = 94-96), (**B**) 1000 nM PCB153 (*n* = 89-91) excluding gastrulation (24-120 hpf), (**C**) L-DOPA 1 mM (*n* = 67-68) or Quinpirole 10 µM (*n* = 120-124) and 100 nM PCB153 alone (*n* = 89-91) and with co-exposure to L-DOPA (*n* = 144) throughout development (4-120 hpf), (**D**) neurotransmitter modulators (L-DOPA *n* = 95-96, L-tyrosine *n* = 92-94, 5-HT *n* = 32, 5-HTP *n* = 78-80, Haloperidol *n* = 175-188, Quinpirole *n* = 106-112, Bicuculline *n* = 161-172, Nipecotic Acid *n* = 140-160) for 30 min in DMSO treated larvae at 6 dpf, (**E**) neurotransmitter modulators (Haloperidol *n* = 91-97, Bicuculline *n* = 142-149, Nipecotic Acid *n* = 169-174) for 30 min in 100 nM PCB153 treated larvae at 6 dpf and 100 nM PCB153 only (*n* = 60-63). Values are presented as mean ± SEM. All data points are biologically independent replicates from at least three independent experiments, *n* varies between stimulus intensities due to varying successful tracking numbers per video. Hollow symbols indicate a significant difference to respective controls (p < 0.01).

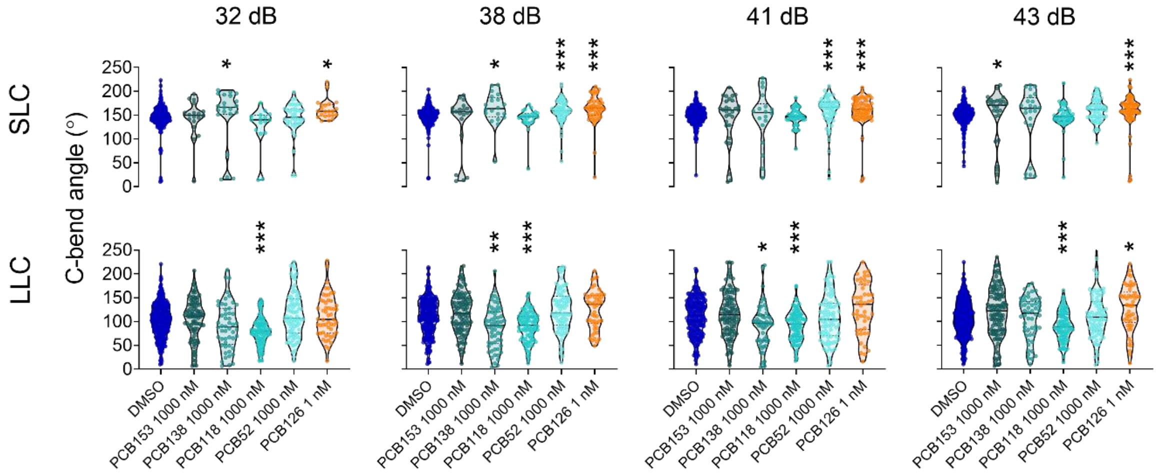


**Figure S6. Bend angle of C-start.** Larval C-bend angle of control larvae remain relatively constant at different auditory stimulus intensities but vary between short latency C-bend (SLC; *n* = 356, median = 152.1°) and long latency C-bend (LLC; *n* = 225, median = 107.9°). PCB153 (1000 nM) exposure resulted in an increased SLC bend angle at the highest stimulus intensity for those few larvae that exhibited an SLC (*n* = 26 of 138, 43 dB, *p* = .0456) while PCB52-treated larvae had increased SLC bend angles (*p* < .001) at two intermediate stimulus intensities (38 and 41 dB). Across all stimulus intensities, PCB126 treated larvae show a significantly increased bend angle during SLC start and PCB118 treated larvae exhibit a decreased bend angle during LLC start in comparison to DMSO treated larvae of respective stimulus intensity. Data points represent biologically independent replicates from at least three independent experiments with black horizontal bars indicating the median. Asterisks indicate significant differences to controls (**p* < 0.05, ***p* < 0.01, and ****p* < 0.001).

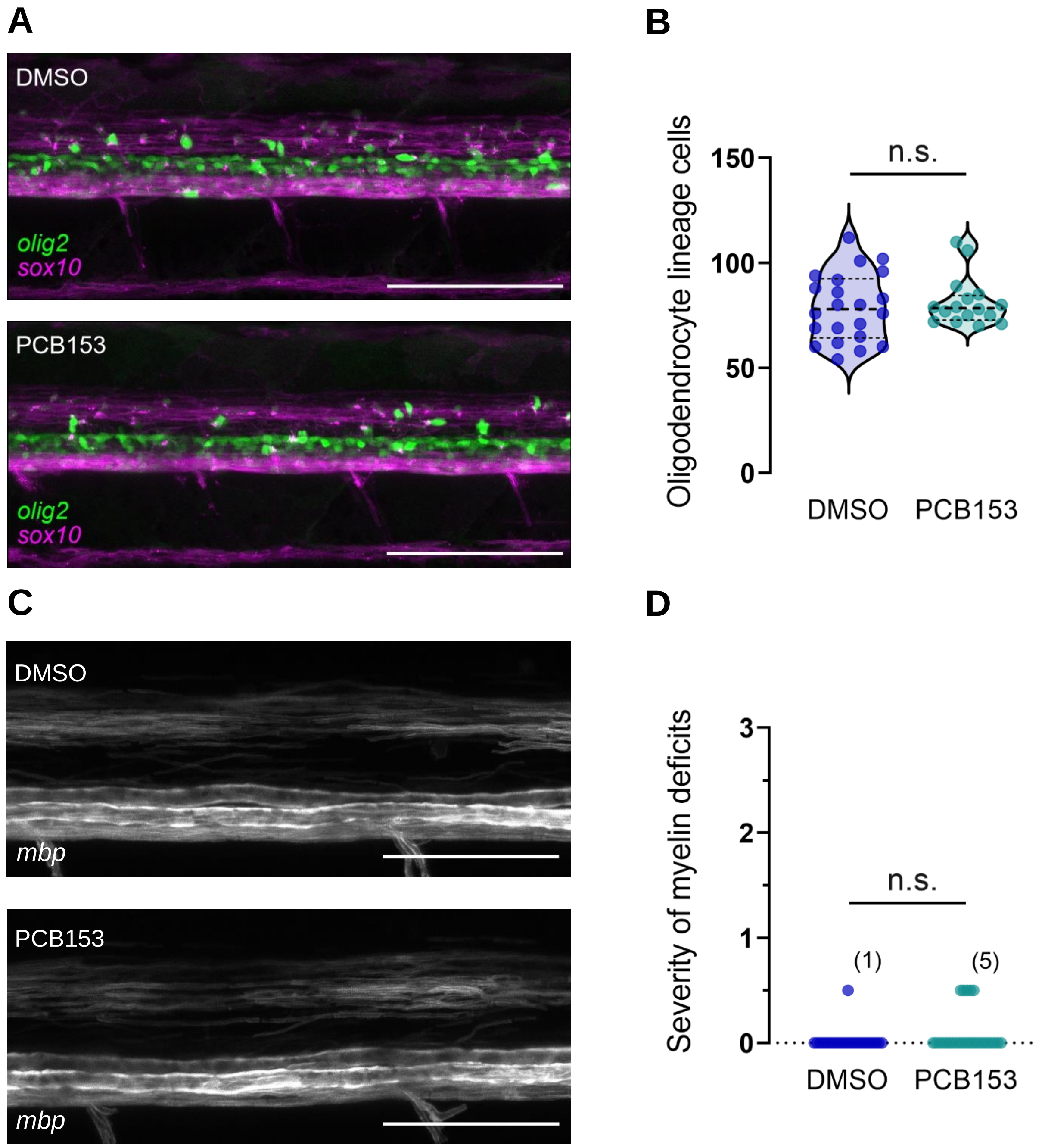


**Figure S7. Myelination in the spinal cord.** (**A**) Representative images of oligodendrocyte precursor cells (OPC) and mature oligodendrocytes (green) in the spinal cords of DMSO (*n* = 22) and 1000 nM PCB153 (*n* = 16) treated *Tg(olig2:EGFP) x Tg(sox10:mRFP)* larvae at 6 dpf. Scale bar = 100 µm. (**B**) Quantification of oligodendrocyte lineage cells (green) per biological replicate in within the same imaging area (319 µm x 160 µm) anterior of the spinal cord across. Data points are biologically independent replicates from three independent experiments. (**C**) Representative images of myelin sheath in the spinal cords of of DMSO and 1000 nM PCB153 treated *Tg(mbp:EGFP-CAAX)* larvae at 6 dpf. Scale bar = 50 µm. (**D**) Classification of severity in myelin deficits in DMSO (*n* = 26) and 1000 nM PCB153 (*n* = 27) treated larvae according to Panlilio et al. 2020. Data points are biologically independent replicates from three independent experiments.

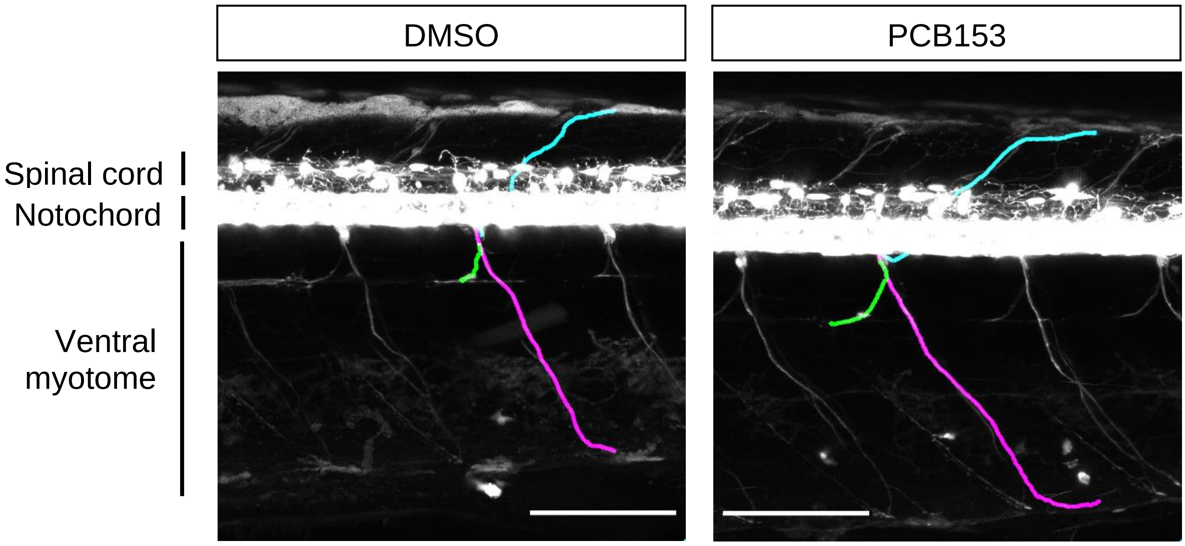


**Figure S8. Morphology of spinal cord primary motoneuron.** Representative images of motoneuron morphology showing no apparent structural deficits in 1000 nM PCB153 exposed larvae (*n* = 18) in comparison to DMSO (*n* = 21). *Tg(olig2:EGFP)* zebrafish were imaged at 6 dpf. The different axons are color labeled as caudal primary (CaP, magenta), middle primary (MiP, cyan), and rostral primary (RoP, green) motoneuron. Scale bar 100 µm.

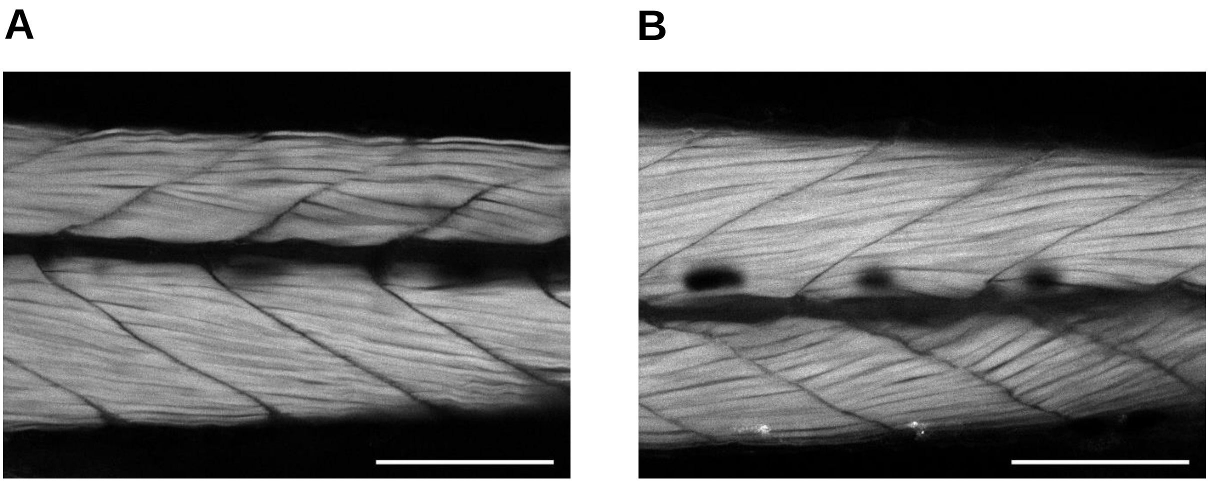


**Figure S9. Fast-twitch skeletal muscles.** Representative images for (**A**) DMSO (*n* = 20) and (**B**) 1000 nM PCB153 (*n* = 20) treated larvae of trunk muscle fibers visualized using labelled phalloidin staining for polymerized actin. Images represent a slice of 1.2 µm thickness out of a z-stack. Scale bar: 100 µm.

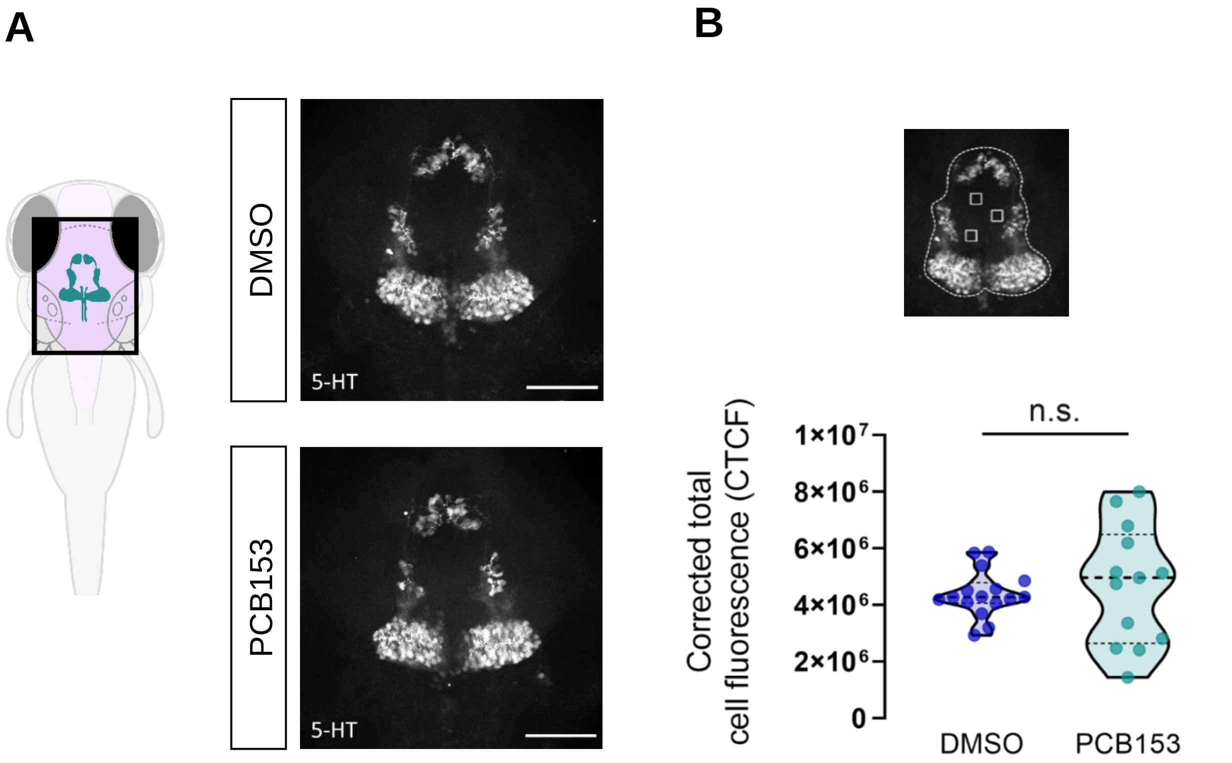


**Figure S10. Serotonin expression in larval zebrafish brain.** (**A**) Diagram indicating serotonergic neurons in the brain and representative images of serotonergic neurons in brains of DMSO ($n$ = 16) and 1000 nM PCB153 ($n$ = 13) treated wild-type (AB) larvae at 6 dpf. Dissected brains were immunostained with anti-5-HT. Scale bar = 50 µm. (**B**) Quantification of 5-HT immunoreactive cell fluorescence calculated using the formula: Integrated density - (ROI x Mean fluorescence of background readings). Dashed line denotes the ROI and the solid square denotes background readings. All data points are biologically independent replicates from three independent experiments.

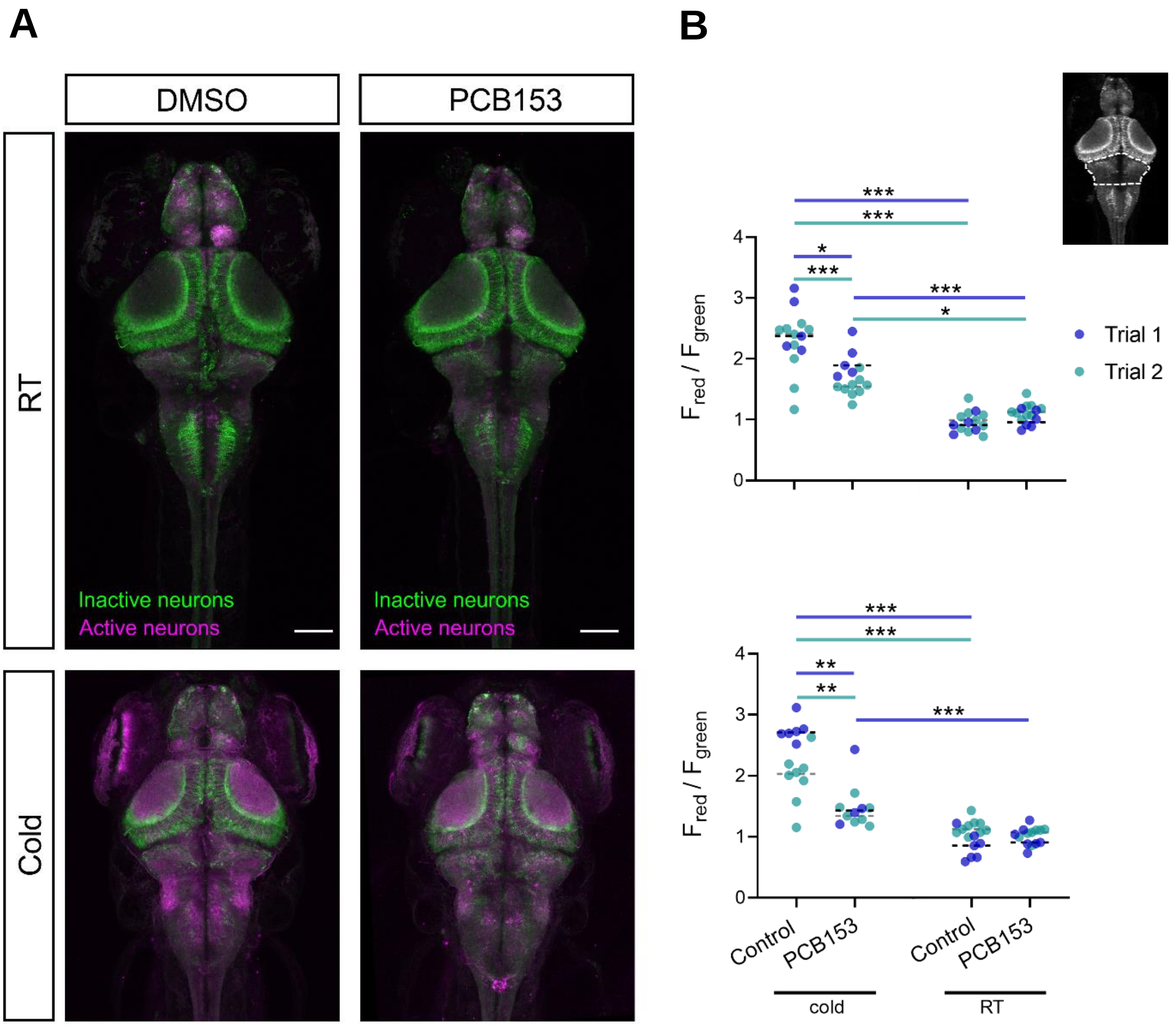


**Figure S11. Active neuronal populations in the brain.** (**A**) Representative maximum intensity z-projection from confocal stack of freely swimming 6 dpf DMSO (*n* = 15) and 1000 nM PCB153 (*n* = 15) exposed CaMPARI larvae at room temperature (RT). Active neurons had elevated calcium levels, which is photoconverted from green to red during UV exposure. Scale bar = 100 µm. (**B**) Ratio of red to green fluorescence intensity in the rostral hindbrain (indicated by dashed line in image) in DMSO and 1000 nM PCB153 exposed freely swimming larvae (upper panel) and startling larvae (lower panel) at room temperature (RT) and in 4 ˚C (cold) medium. Asterisks indicate significant differences to controls (*p < 0.05, **p < 0.01, and ***p < 0.001).

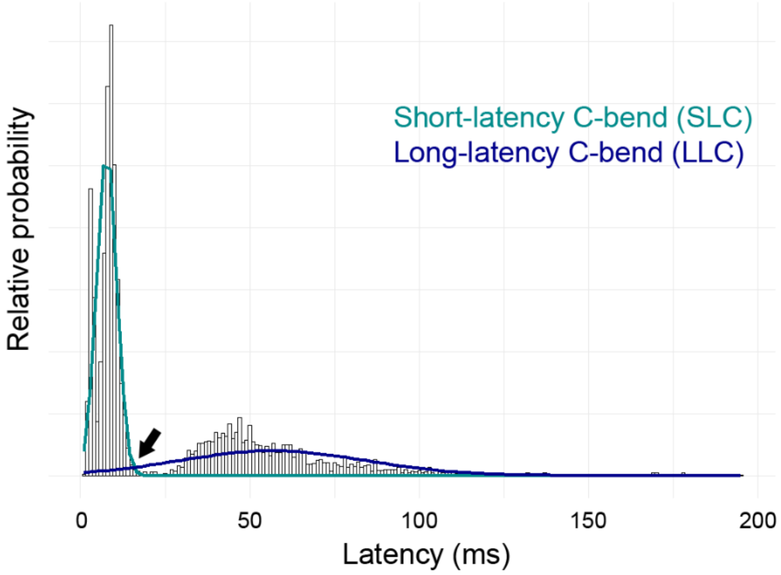


**Figure S12. Histogram of the latency distribution for control larvae.** A two-component Gaussian mixture model was run to determine the cut-off by which there is a greater than 50% probability of a given data point belonging to either modeled distribution. The cut-off value is 15 milliseconds and is indicated by a solid arrow.
